# Spatial engineering of *E. coli* with addressable phase-separated RNAs

**DOI:** 10.1101/2020.07.02.182527

**Authors:** Haotian Guo, Joseph C. Ryan, Adeline Mallet, Xiaohu Song, Victor Pabst, Antoine Decrulle, Ariel B. Lindner

## Abstract

Biochemical processes often require spatial regulation and specific microenvironments. The general lack of organelles in bacteria limits the potential of bioengineering complex intracellular reactions. Here we demonstrate Transcriptionally Engineered Addressable RNA Solvent droplets (TEARS) as synthetic microdomains within the *Escherichia coli.* TEARS are assembled from RNA-binding protein recruitment domains fused to poly-CAG repeats that spontaneously drive liquid-liquid phase separation from the bulk cytoplasm. Targeting TEARS with fluorescent proteins revealed multilayered structures and a non-equilibrium mechanism controlling their composition and reaction robustness. We show that TEARS provide organelle-like bioprocess isolation for sequestering biochemical pathways, controlling metabolic branch points, buffering mRNA translation rates and scaffolding protein-protein interactions. TEARS are a simple and versatile tool for spatially controlling *E. coli* biochemistry.

## Introduction

Living systems coordinate complex biochemical reactions using intracellular spatial organization, a powerful strategy that bioengineers have long sought to replicate. In eukaryotes, membrane-bound organelles create an isolated environment (*1*) that can be reprogrammed by fusing components of interest to organelle-targeting domains (*2*). However, prokaryotes generally lack universal organelles, posing an attractive challenge to engineer synthetic alternatives. To this end, modular DNA-, RNA- and protein-based scaffolding, widely found in natural systems (*3, 4*) were engineered (*5*) or rationally designed (*6, 7*) for various metabolic engineering applications (*5–7*) and control of gene expression (*8, 9*). Engineering such scaffolds is hindered by difficulties to cluster large numbers of different enzymes (*7*), or to delocalize enzymes to prevent unwanted crosstalk (*10*).

In this study, we chose to couple mimicry of eukaryotic membraneless organelles (*11*) with protein targeting to provide recruitment specificity. Membraneless organelles are formed through a common liquid-liquid phase separation (LLPS) process that condenses a variety of biopolymers (*11, 12*). Biopolymer condensates can be derived from a single component (*12, 13*) and may adopt flexible sizes or shapes (*11*), facilitating rational design. In comparison, membrane-bound organelles require complex processes of biogenesis and regulation (*14*), and protein-shelled microcompartments have precise multicomponent assembly of fixed shapes and sizes (*15*).

Previous works of artificial LLPS focused mainly on protein-driven systems, based on translational fusion of an LLPS-domain with effector proteins *in vitro* (*16*) or in eukaryotic cells (*17, 18*). Though successfully demonstrating the validity of the approach, these studies revealed limitations that hamper generalization and scaling, including reduced activity of fusion enzymes, and requirements for specific expression tuning of the assembling proteins (*12, 17*). In addition, many of those proteins are highly aggregation-prone in bacteria (*19*).

In an attempt to overcome these hurdles, we introduce here the first RNA-based modular system for synthetic LLPS, Transcriptionally Engineered Addressable RNA Solvent droplets (TEARS). TEARS feature a transcriptional fusion of an LLPS domain to an aptamer-based protein recruitment domain, forming an intracellular separated *RNA solvent*, to which additional components (*solutes*) may be scaffolded to introduce customized functionalities (Fig. 1A). Phase separation is achieved using CAG triple-ribonucleotide (rCAG) repeats. Though RNAs are generally considered as regulators of LLPS (*20*), the triplet repeat expansion is known to trigger liquid-like pathological puncta in human cells (*21*). However it was unclear whether relevant proteins such as muscleblind-like-1 (MBNL1) (*21*) were required for LLPS. Here we consolidate our preliminary results by demonstrating that such rCAG repeats are sufficient for condensate formation in the absence of any other human cytosolic factors (*22*) and provide evidence for their liquid-like properties and reversible assembly dynamics (*11–13*) of the rCAG solvents.

**Fig. 1.**
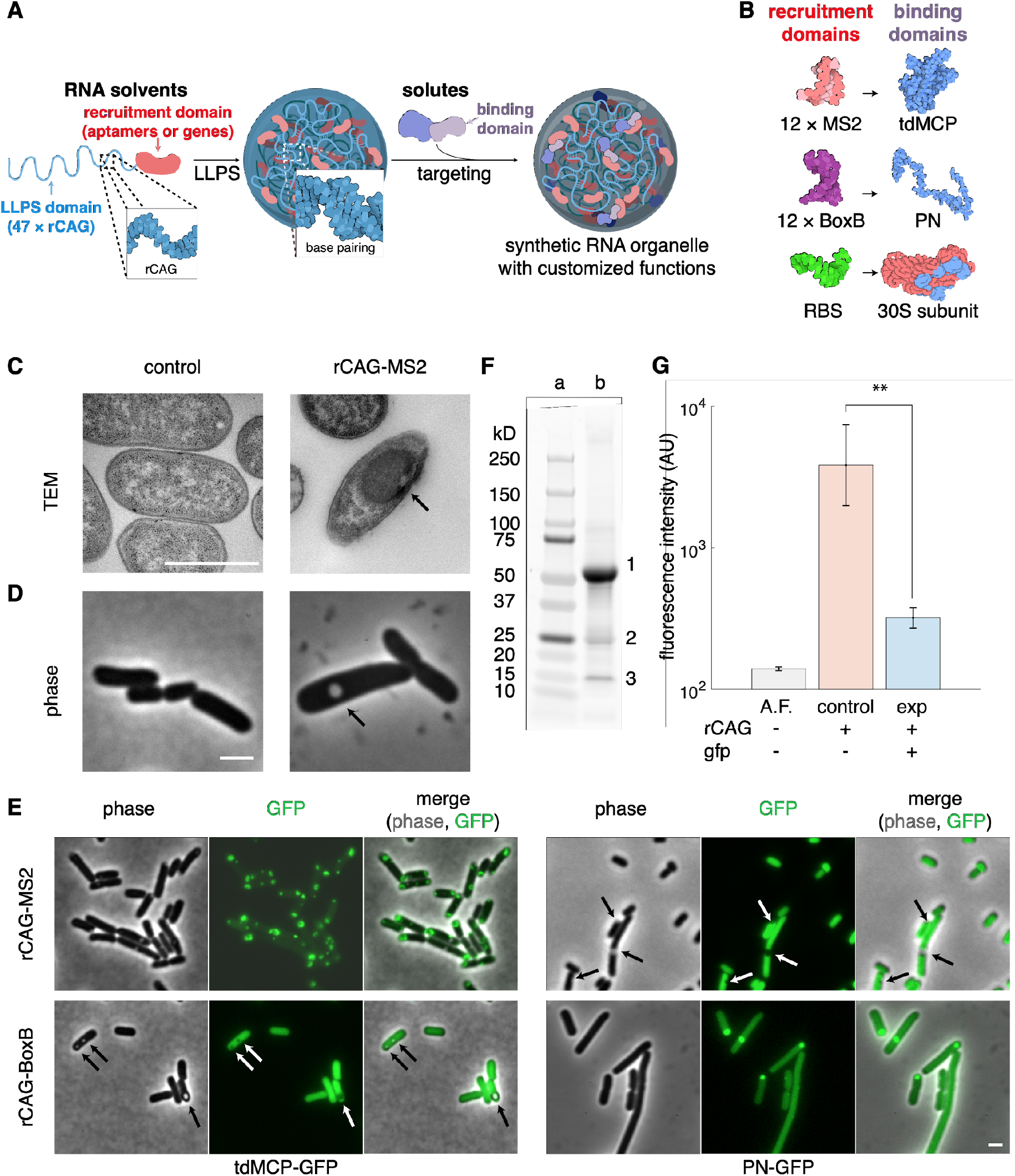
Design, construction and identification of TEARS with selective permeability. (A) Schematic representation of TEARS. TEARS are composed of TEARS and recruited solutes. TEARS consists of multivalent base-pairing LLPS RNA domains fused to a recruitment domain made of aptamers or translational units of genes. Solutes can be targeted into TEARS, through binding to either recruitment domains (shown in the diagram), or other recruited solutes (not shown). (B) Recruitment systems: (CAG)_47_ LLPS domain was fused to (MS2)_12_, (BoxB)_12_ aptamer recruitment domains, or translational units of sfGFP expression (rCAG-MS2, rCAG-BoxB, and rCAG-RBS-gfp), which interact with tdMCP- or PN-tagged proteins (e.g. GFP), or 30S ribosomal subunit, respectively. (C and D) Synthetic RNA condensates: TEM imaging (C) and phase contrast images (D) of control cells (left) and cells expressing rCAG-MS2 (right) under a tetracycline-inducible promoter, following overnight induction by 50 ng/ml anhydrotetracycline (aTc). Arrows highlight the highly-dense assembly. (E) LLPS of CAG-aptamer is independent from RBD-GFP binding, judged by phase contrast and fluorescent imaging of cells induced for 2 hours in exponential phase. Shown from top to bottom are rCAG-MS2, rCAG-BoxB, and from left to right are tdMCP-GFP, and PN-GFP. In each indicated system, shown from left to right are phase contrast, GFP, and overlay images of representative cells. Arrows highlight representative foci void of fluorescence. Scale bars, 1 μm. (F) Sodium dodecyl sulfate–polyacrylamide gel electrophoresis (SDS-PAGE) of condensate purified from BL21AI cells expressing both tdMCP-GFP and rCAG-MS2. (a) protein ladders (molecular mass shown on the left), (b) purified condensate. Highlighted bands are (1) complete tdMCP-GFP, (2) tandem dimeric, and (3) monomeric fragments of MCP, judged by molecular mass. (G) rCAG-mRNA sequesters translations for 14-fold. Shown from left to right are constructs to measure autofluorescence (A.F., grey), rCAG-RBS-gfp (blue) or RBS-gfp (red). and flow cytometry measurement of median fluorescence intensity in induced cells (right). AU, arbitrary units. **, p < 0.01, in paired Student’s t-test.

TEARS are shown to exhibit programmable selective permeability, acting as solvents only for specific aptamer-targeted components. Endogenous cytosolic proteins are effectively excluded, creating an isolated environment that we harness here for versatile fundamental and synthetic applications. The modular approach herein facilitates the study of protein-targeting driven phenomena that were seldom addressed (*11–13*), and of non-equilibrium tuning, mostly unattainable in vitro (*13*). In particular, we provide a simple, alternative mechanism for multiphase demixing (*16, 23*) and compositional control (*24*), and unveil diverse regulation of dynamic robustness beyond noise buffering (*25*). We then explore potential biotechnological applications, in buffering mRNA translation to generate constant expression independent of ribosomal binding sites (RBS) strength, in switch-like shunting metabolic flux of deoxyviolacein biosynthesis (*26*), in sequestering lycopene biosynthesis by relocating membrane-associated enzymes (*27*), and in clustering of α-complementation of endogenous β-galactosidase mutants (*28*) for growth rescue.

## Results

### Ribonucleic CAG-repeats form RNA condensates with selective permeability

Transcription of 47x repeats of rCAG followed by 12x repeats of the MS2 aptamer (rCAG-MS2) (Fig. 1B) in *E. coli* produced highly dense cytosolic assemblies as judged by transmission electron microscope (TEM) and phase contrast (Fig. 1, C and D). Condensate formation was dependent on rCAG-repeat transcription but independent of which aptamer was used, implying that rCAG alone drives RNA condensation. Expression of green fluorescent protein (GFP) tagged with the Aptamer-Binding Domain (ABD), tandem-dimeric MS2 coat protein (tdMCP-GFP), resulted in fluorescent RNA condensates (Fig. 1E, S1A). The tdMCP-GFP labeling was robust to temperature and bacterial background strain (Fig. S2). Extending our toolbox by modular replacement of the aptamer domain with BoxB and the ABD with Protein N (PN) produced similar rCAG-BoxB condensates with PN-GFP labels (Fig. 1B). Importantly, unrecruited proteins were generally excluded from the RNA condensates as non-cognate aptamer-ABD combinations showed no specific labeling as visible fluorescence voids overlapping the formed phase-bright RNA untargeted condensates (Fig. 1E, S1B) were observed. These results demonstrate that TEARS exhibit selective permeability, discriminating solute targeted proteins from untargeted cytosolic proteins. Further validation of this result was obtained using denaturing electrophoresis of condensates purified from cells expressing rCAG-MS2 (Figs. 1F, and S3). The ability of TEARS to exclude bulk cytoplasm may explain their limited toxicity and minimal effects on cellular growth (Fig. S4).

In line with our conclusion that transcription of CAG triplet repeats drives the observed condensation, we tested the effect of fusing rCAG repeats to a translational unit overexpressing superfolder GFP (rCAG-RBS-gfp). Measured by flow cytometry, rCAG-RBS-gfp expressed 14-fold lower GFP compared to an untagged RBS-gfp mRNA (Fig. 1G), indicating sequestration of mRNA from the translation machinery. Taken together, the RNA condensates are capable of programmable, selective recruitment and exclusion of additional components, and thus can be considered as spatially distinct compartments, reminiscent of membraneless organelles (*11*).

### Liquid-like properties of TEARS inside bacterial cells

Next, we set to portray key physical properties of TEARS. TEARS that contact cytoplasmic membranes often display concave meniscus (Fig. 2A and B), as characteristic surfaces. This indicates the existence of both cohesion and adhesion, which distinguishes liquids from solids (*29*).

**Fig. 2.**
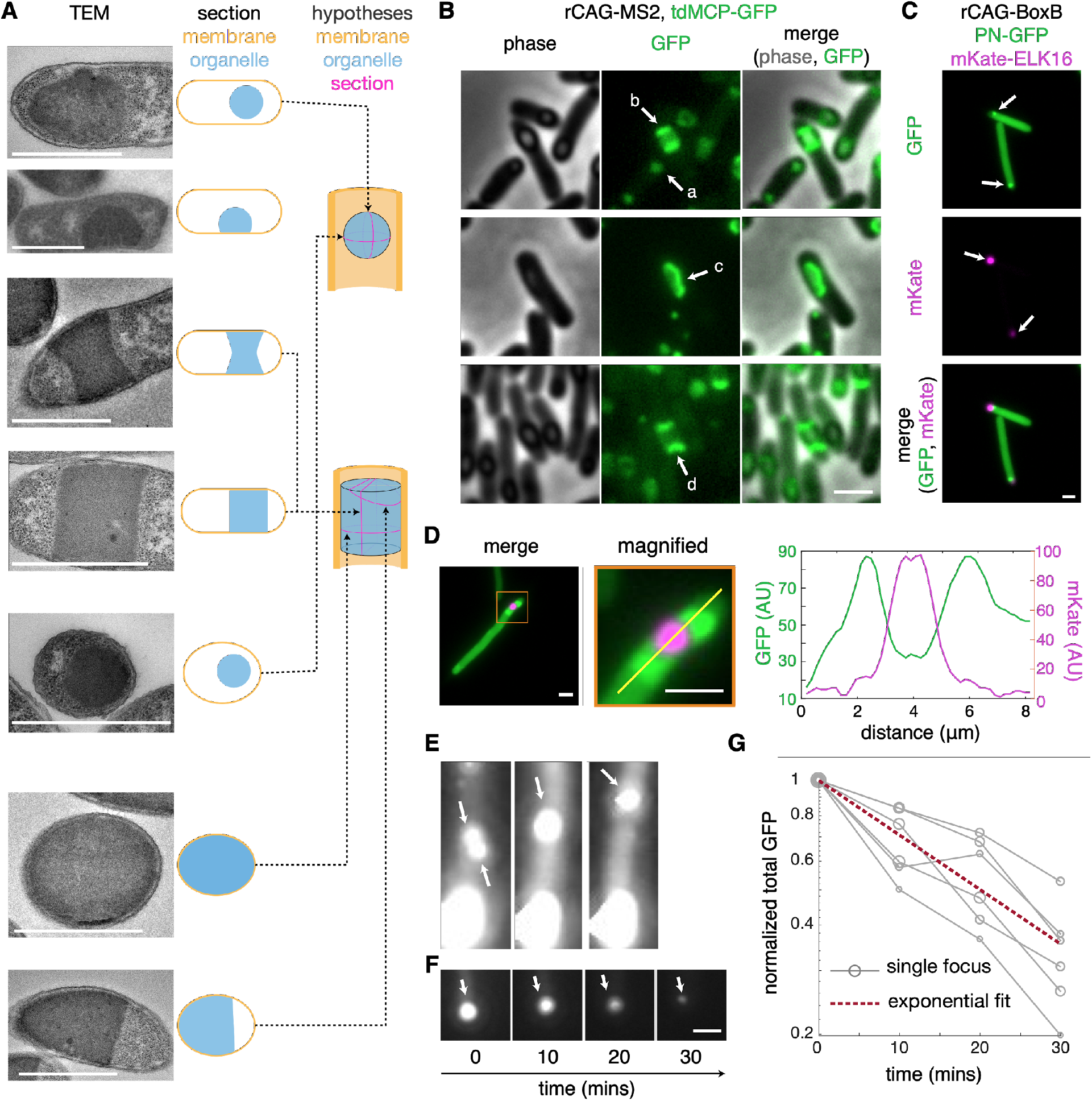
Liquid-like shapes and dynamics of TEARS. (A) TEM images and schematic diagrams of synthetic RNA foci sections. Shown from top to bottom are examples of sphere, hemisphere, longitudinal section of hyperbolic and right cylinder with concave or flat meniscus, cross section of cylinder, and intersection of cylinder that shows half ellipse, as represented in adjacent color-coded schemes. 3-dimensional shapes of spherical droplet and cylinder, with arrow links and magenta lines, indicates the respective sections shown in TEM. (B) Phase contrast (left) and GFP images (right) of identified different shapes of the synthetic RNA foci, highlighted by arrows, (a) sphere, (b) flat meniscus, (c) hemisphere, and (d) concave meniscus. (C) Fluorescence images of cells coexpressing mKate-ELK16, rCAG-BoxB and PN-GFP. Shown from top to bottom, GFP, mKate, and overlay, with arrows highlighting RNA and protein foci. (D) Results of random collision between RNA foci and protein aggregates. Shown from left to right, overlay, magnified image of the representative collision (orange box), and line profile for the GFP and mKate channels (yellow line, green and magenta profiles, respectively). AU, arbitrary units. (E) Time-lapse snapshots of fusion between RNA droplets in cells expressing rCAG-MS2 and tdMCP-GFP. Droplets are highlighted before and after fusion by arrows. (F) Progressive dissolving of a representative RNA droplet (highlighted by arrows) during cell growth. (G) Fluorescence loss as an exponential function of time. Total fluorescence is calculated as the sum of fluorescence intensity in the segmented area, and then normalized by the initial value at time 0. Each grey line shows fluorescence intensity of a representative focus, and circles represents the focus sizes. Red dashed line fits the data of 6 focus, with half-life time ~ 20 minutes, 95% confidence bounds as 17-25 minutes (R^2^ = 0.823). Scale bars, 1 μm.

As liquids, the shape of TEARS is governed by surface tensions and objects they contact, most likely forming spherical droplets to minimize surface area, or adapting to the shape of cytoplasmic membranes (Fig. 2A and B). Occasionally, TEARS adapt irregular shapes when in contact with structures as the nucleoid (Fig. S5A) or the cellular division ring (Fig. S5B). We observed that cellular diameter increased when TEARS tethered to the cytosolic membrane at polar positions (Fig. S6), indicating strong pressure on the cell wall resulting from surface tensions or other perturbation of cell-wall synthesis. We further challenged TEARS’ shape by co-expressing synthetic protein assemblies driven by ELK16 aggregating peptide (*30*) fused to a red fluorescent protein (mKate) (mKate-ELK16). Coexpression of mKate-ELK16, rCAG-BoxB and PN-GFP produced non-overlapping, coexisting RNA and protein foci (Fig. 2C). Random collisions immersed protein aggregates within the GFP-labeled TEARS (Fig. 2D), proving the latter’s liquid-like shape adaptation.

In order to demonstrate liquid-like dynamics in real time, we placed cells in the mother-machine microfluidics setup (*31*), briefly pulsed the expression rCAG-MS2 and tdMCP-GFP to form fluorescent droplets, and then chased their behaviors (*32*). We documented droplet fusions (Fig. 2E), a hallmark of fluidity (*12*). In cases of coexistence of multiple droplets within a cell, smaller ones disappeared as the size of larger ones increased, suggestive of Ostwald ripening (Fig. S7) (*23*). Without sufficient induction, the total fluorescence of the droplets decreases exponentially with a mean half-life of 20±4 minutes, regardless of the initial sizes (Fig. 2F and G), suggesting dilution-driven dissolution of TEARS as a result of bacterial exponential growth.

### Coexisting multiple phases underlying single-solute enrichment

Engineering of TEARS may facilitate understanding of LLPS physics. Complementing recent studies that focused on LLPS proteins (*11–13, 19*), we adopt a solute-centered view of phase separation. We took advantage of the ABD-tagged fluorescent protein reporters, to study recruitment and partitioning dynamics between the cytosol and TEARS. We first characterized the simplest case where the binding between a single solute and its recruitment domain dominates the behavior of the system (Fig. 3A). Co-induction of rCAG-aptamers to govern the spatial distribution of the ABD-GFP solutes, resulted in strong partitioning into RNA over the aqueous phase, leading to a 2.6- or 1.3-fold fluorescence enrichment within TEARS, for MS2- and BoxB-based systems, respectively (Fig. 3B).

**Fig. 3.**
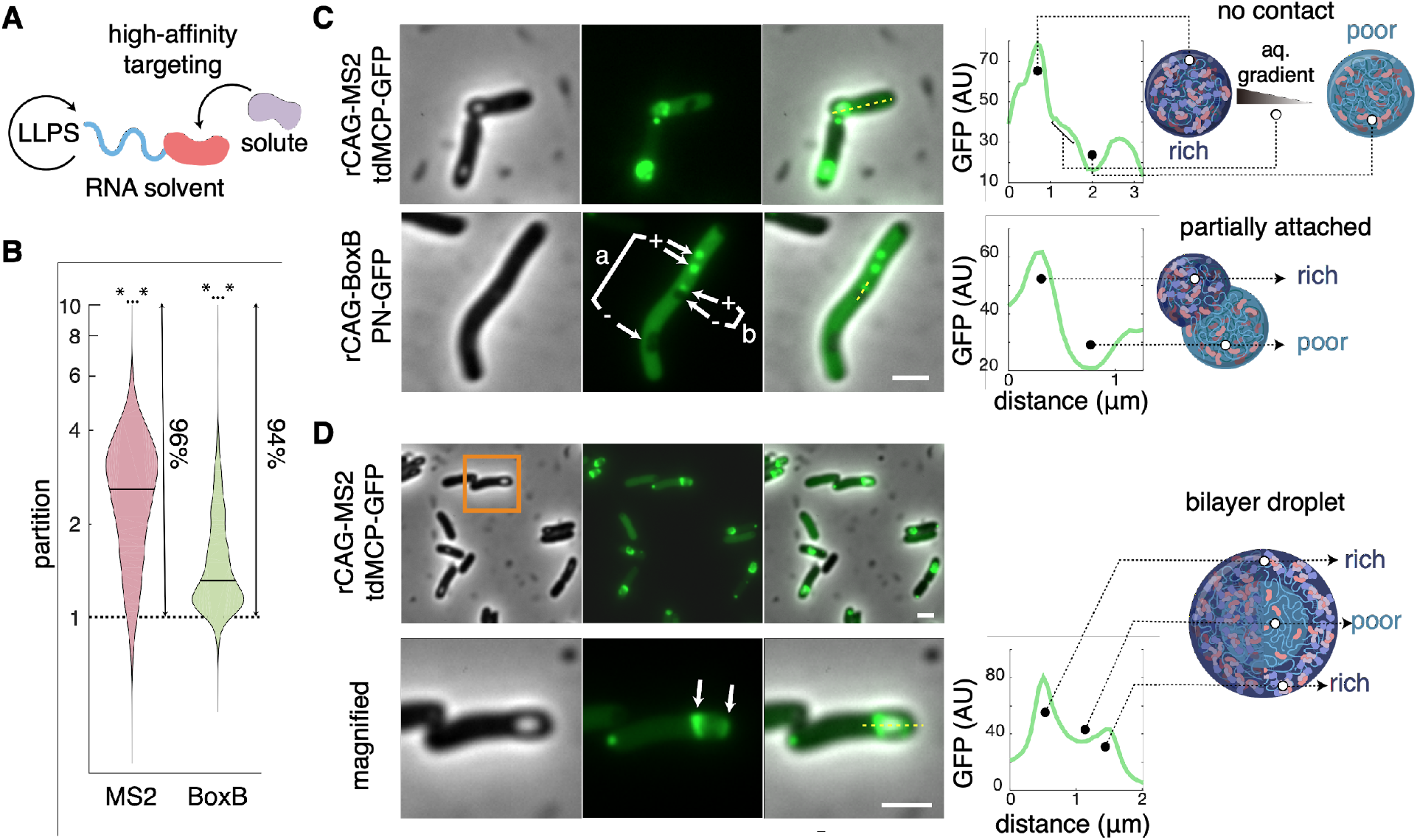
Multiphase separation and multilayer architectures of TEARS. (A) Schematic of the single-solute targeting through high-affinity binding. (B) Quantification of partition, measured by the ratio of mean fluorescence intensities of each identified focus over respective cytosol. From left to right, violin plots displayed distributions of measurements for 13646 and 4109 foci identified in 6682 and 995 cells expressing MS2 and BoxB based systems, respectively. *…*, significantly greater than 1, p value < 2^−1022^, in one-tailed z-test on logarithmic scale. Arrows highlighted the range greater than 1, and percentages. (C) Representative coexistence of foci (+) enriched GFP and (-) void of GFP, either with (a) no contact or (b) shared interface, in cells expressing MS2-based (top) or BoxB-based (bottom) systems. Shown from left to right are phase contrast, GFP, overlay, line profiles for GFP channel (yellow dashed lines, GFP profiles), and diagrams of profiled foci (dashed lines linking region in profiles and respective phases in diagrams, rich, GFP-rich phase, and poor, GFP-poor phase). Shown on the top profile, a flatter gradient exists in the aqueous cytosolic region (aq.) between GFP-rich and GFP-poor foci; shown at the bottom, smooth transition from GFP-rich to GFP-poor phase implies a shared interface between partially attached foci. AU, arbitrary units. (D) Many MS2-based foci only enriched GFP in the outer layer. Shown from left to right are phase contrasts, GFP (arrows highlighted GFP-rich sides), overlay, line profiles for GFP channel (yellow dashed lines, green profiles), and schematic diagram of bilayer two-phase droplets (dashed lines linking region in profiles and phases in diagrams, rich, GFP-rich phase, and poor, GFP-poor phase).

Intriguingly, single-solute enrichment further induces the coexistence of multiple phases. Under the condition of rCAG-aptamer overexpression, we found cases of GFP-rich and GFP-void droplets coexisting in the same cells (Fig. 3C), indicating distinct two phases at a quasi-static non-equilibrium state. Mixing may not occur even when GFP-rich and -free droplets are partially attached (Fig. 3C). Moreover, even though all of the large rCAG-MS2 droplets are filled by RNAs as judged by TEM images (Fig. S8A), a majority of them displayed tdMCP-GFP only at the cytosol-contact surfaces (Fig. 3D, and S8B). This suggested the establishment of a metastable bilayer, consisting of RNA-only internal layer and RNA-protein outer layer. A simpler explanation that the core of TEARS are less accessible, is ruled out given the formation of fully-fluorescent droplets with micron-diameter sizes by pulse induction (Fig. 2E and F).

This observation is akin to published theoretical models that simulate LLPS droplets demixing, triggered by bivalent protein association (*34*). Thus, coexisting phases and multilayered architectures can emerge from a simple LLPS system driven by the interaction between one phase-separating molecule (rCAG-aptamer) and a single solute (ABD-GFP; Fig. 3A). In comparison, previous multilayered systems required at least two immiscible LLPS proteins involving multiple domains (*16, 34*). From an application perspective, multiphase separation stabilizes local enrichment of solutes of interest under excess of RNA solvents. This is in contrast with previously reported scaffolding where overabundant scaffold proteins titrate targeted components away from each other (*4, 5*).

### Modulating TEARS composition and bimolecular binding efficiencies by a non-equilibrium mechanism

We next studied the partitioning of solutes between the cytosol and the TEARS at non-equilibrium, cellular growth conditions where the binding processes may not be dominant. How would the localization of targeted solutes and the efficiency of their binding reactions be affected by solute synthesis and turnover (dilution, degradation, or consumption)? To enable comparison between condensate-organized and cytosolic well-mixed reactions (*e.g.,* in absence of condensates), we considered a modular framework where condensates directly recruit a primary molecule (*receptor*), that in turn may recruit a secondary solute (*ligand*) to the condensates through receptor-ligand association (Fig. 4A). We termed this process as “ligand sorting” to signify selectivity conferred by such specific interactions.

**Fig. 4.**
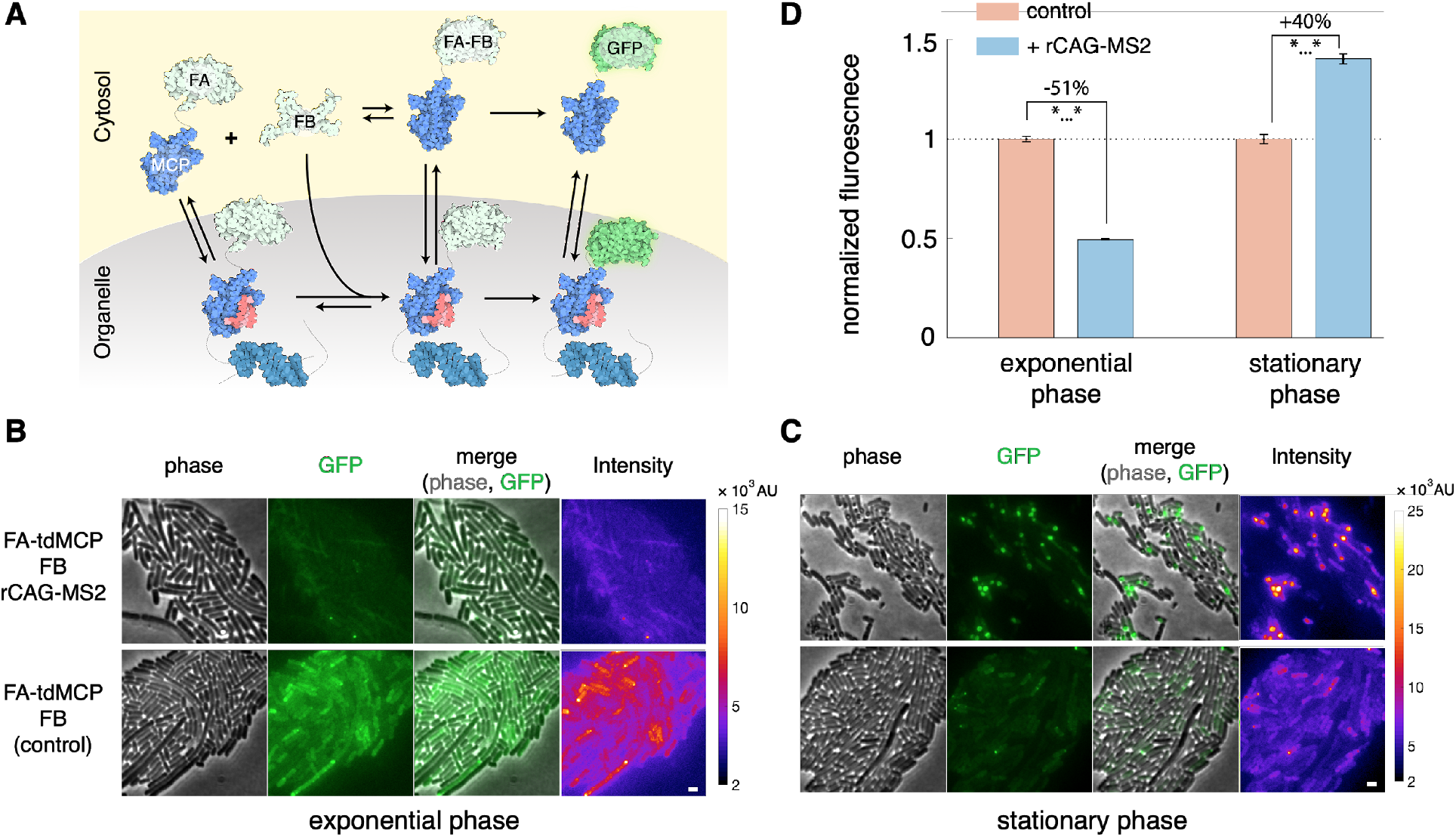
Ligand sorting as a simple mechanism of compositional controls and reaction tuning. (A) Schematic representation of the BiFC system used for experimental validations. FA-tdMCP and FB can bind to each other, form FA-FB complexes, and then irreversibly mature into GFP. tdMCP-tagged FA, FA-FB complexes, and GFPs are partitioned between TEARS and cytosol through high affinity binding to the MS2 aptamers (red). FB may enter droplets through FA-FB association. FA and FB were continuously synthesized. All components were diluted or degraded in cytosol or in droplets. (B and C) Images of representative fully-induced cells with indicated systems under rapid growth in exponential phase (B) or growth arrest in the stationary phase (C). Shown from left to right are phase contrast, GFP, overlay, and fluorescence intensity heatmap images. Scale bar, 1 μm. Colorbars show the fluorescence intensity. A.U., arbitrary units. (D) Quantification of normalized whole-cell fluorescence intensity per pixel, cultured either in exponential (left) or stationary phases (right). Fluorescence intensities are normalized by the median value among cells expressing FA-tdMCP and FB alone. Errors are standard deviations of the mean fluorescence calculated with bootstrapping. *…*, p-values < 10^−15^, by pairwise Kolmogorov–Smirnov test.

To this end, we established a reporter assay (*4*), by overexpressing the tdMCP-tagged one half of split GFP (FA-tdMCP) as the receptor, and the other half (FB) as the ligand (Fig. 4A). We measured receptor-ligand binding by bimolecular fluorescence complementation (BiFC), and explored the effect of turnover rates by contrasting two distinct regimes: exponential growth with fast turnover governed by dilution, and stationary phase with very slow turnover due to proteolytic stability. Under exponential growth, scaffolding of FA-tdMCP to the rCAG-MS2 droplets displayed no GFP enrichment (Fig. 4B) and reduced whole-cell fluorescence by 51% compared to expressing FA-tdMCP and FB alone (Fig. 4D). This is in stark contrast to abundant fluorescent foci in absence of dilution (*e.g.,* in stationary-phase state) (Fig. 4C), where whole-cell fluorescence increased by 40% (Fig. 4D), probably due to reduced GFP degradation or enhanced FA-FB assembly by rCAG-MS2 scaffolding. Some foci exhibited signature multilayered structures that differed themselves from aggregates (Fig. S9), supporting enrichment of completed GFP by the liquid-phase TEARS.

Taken together, TEARS resemble natural multicomponent organelles, seeming to control composition and reactions by non-equilibrium dynamics. Particularly, compositional control through ligand sorting may only require one pair of interacting molecules, where TEARS with the same binding sites (FA) can display different preferences to the ligands (FB) under various turnover rates. In comparison, in the equilibrium mechanism known as “scaffold-client” model (*24*), recruitment preference is controlled by the stoichiometry of multiple scaffolds with different binding sites.

### Phase separation provide diverse mechanisms to control reaction robustness

Next, we explored how the TEARS can be used to regulate fluctuations of the model receptor-ligand binding reaction. Many cellular systems are insensitive to reactant supplies, a property known as dynamic robustness of the reaction (*35*). Yet, tuning robustness is key to biological systems to either buffer environmental changes and intrinsic noise (*36*) or to increase variability to overcome hazards by bet-hedging and to drive development and evolution (*37*). Phase separation was previously described as a mechanism to reduce noise by suppressing concentration fluctuations in the aqueous phase against different supplies of LLPS molecules, as shown in vitro and recently in vivo (*11, 25*). Here we extend this simple assertion by illustrating diverse regulations of robustness.

Firstly, given the small number of TEARS per cell (typically ≤ 3), even in absence of an active sorting mechanism (*38*), cellular division will result in asymmetric distribution between daughter cells, which induces distinct phenotypes and increases the variability of cytosolic concentration (Fig. S10). Moreover, we showed that coexistence solute-rich and -poor phases established intracellular asymmetry of cytosolic gradient (Fig. 3C, and S11) that may be inherited by daughter cells. The stability of such quasi-static gradients can be broken after division, causing extra fluctuations. Overall, this results in stronger noise as measured by a bigger coefficient of variations (CV) of ABD-GFP fluorescence in presence of rCAG-aptamer in both cellular or cytoplasmic (e.g. cellular volume excluding the droplets) volumes (Fig. 5A).

**Fig. 5.**
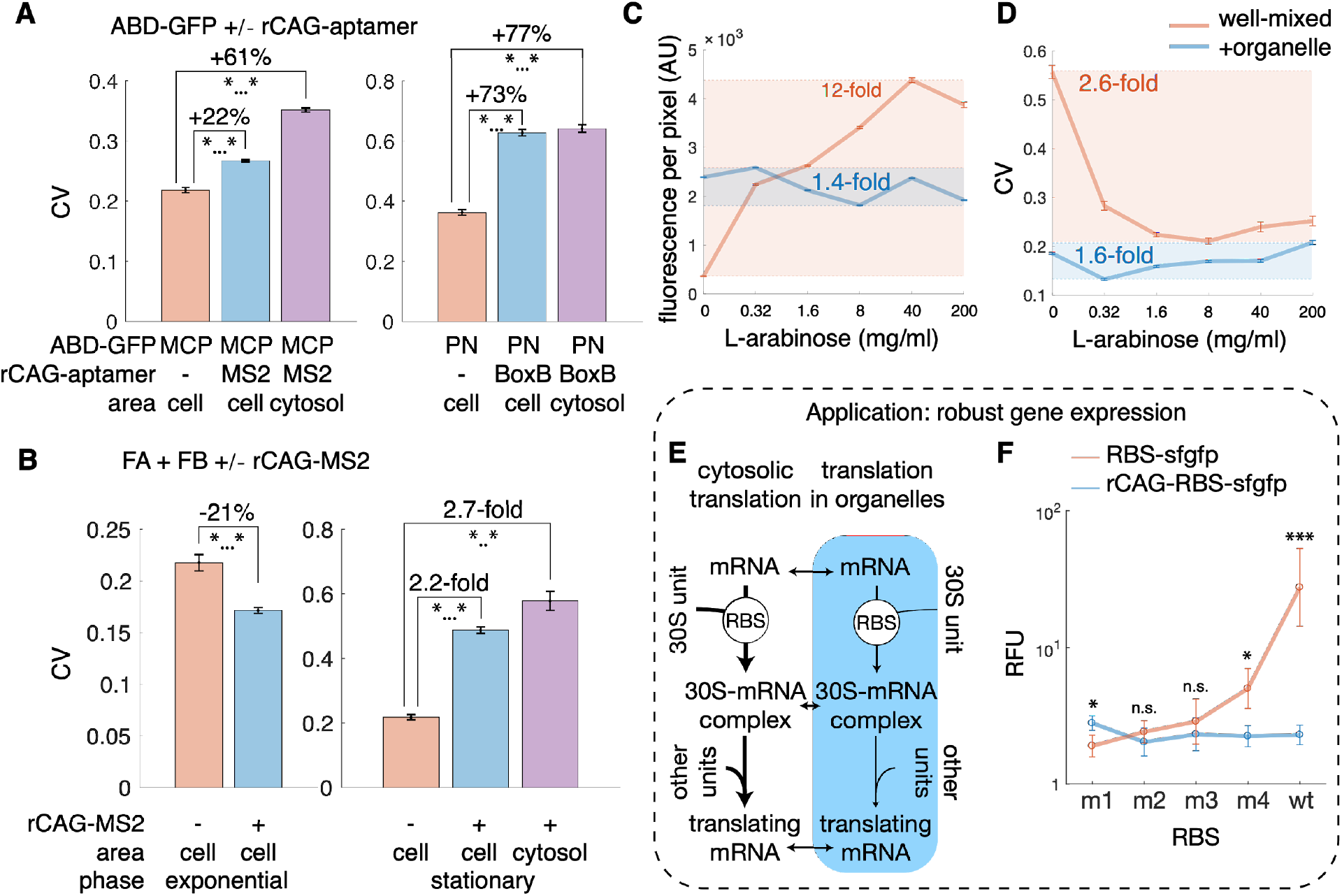
LLPS-driven concentration robustness and cellular homeostasis. (A) Noise strengths of fluorescence per pixel in the whole cell or cytosol in cells expressing indicated ABD-GFP alone or with cognate rCAG-aptamers under the same induction levels. In cells without rCAG-aptamers, cells and cytosols are considered the same. Noise is measured by coefficient of variation (CV), defined as standard deviation divided by mean values. (B) Noise strengths of fluorescence per pixel in indicated areas in cells expressing FA-tdMCP, FB alone or with rCAG-MS2, cultured in exponential (left) or stationary phase (right). *…*, p-value < 10^−70^, by pairwise Kolmogorov– Smirnov test. (C and D) Fluorescence per pixel (E) and noise strengths (F) in exponential-phase cells expressing FA-tdMCP, FB alone (red) or with (blue) rCAG-MS2 expression, under indicated L-arabinose inductions. Errors are standard deviations from bootstrapping. Fold changes are calculated as the ratio between maximal and minimal mean values. AU, arbitrary units. (E and F) Phase separations of mRNA stabilizes translational efficiencies. (E) Dynamics of rCAG-mRNAs translation. mRNA partitions between mRNA droplets (blue) and cytosol. RBS strength governs ribosomal binding in cytosol and droplets. aa-tRNA, aminoacyl-tRNAs. Due to the high turnover rates of translational units in cells, we expected translations in droplets to be significantly suppressed according to the ligand-sorting framework. (F) Flow cytometry measurements of GFP translated from rCAG-RBS-gfp (blue) and RBS-gfp (red) variants. RBS measured in Figure 1H is termed as the wild type (wt), and the rest variants are termed as m1 to m4 according to their strengths. Shown from top to down are distributions displayed as histograms, and average medians. Errors are measured values from 3 biological replicates. RFU, relative fluorescence units compared to autofluorescence. *, p-value < 0.05, ***, p-value < 0.005, n.s., not significant, by pairwise Student’s t-test.

Condensates can be more complex than a single-entity system (*11–13*). The solute partitioned by condensates needs to interact with downstream molecules to output functions. In the presence of heterotypic interactions, solutes may no longer have a fixed concentration in the aqueous phase (*33*), raising the question of whether noise is still buffered in such multicomponent systems. This process can be mimicked in our ligand-sorting framework. By measuring the concentration of ligand-receptor binding products as the output, we found that variability relies on turnover rates as verified by BiFC assays. During exponential growth, despite the above-described effect of asymmetric division contribution to increasing variability, there was an overall 21% reduction of variability due to FA-tdMCP scaffolding by TEARS (Fig. 5B). In contrast, during the stationary phase, droplets sequentially recruited FB (Fig. 4E), resulting in strongly amplified variability for 2-3 fold (Fig. 5B).

Lastly, TEARS may also regulate robustness towards deterministic changes of reactant supplies (*36*). Sensitivity analysis of a simple mathematical model suggests that the more droplets inhibit receptor-ligand binding, the total product concentration is less sensitive to the synthesis rate of ligands and receptors (supplementary text 1.3). This is corroborated experimentally by tuning synthesis rates (Fig. S12). In absence of the rCAG-MS2 droplets, mean fluorescence increased and variability decreased with higher synthesis rate, up to 12-fold and 2.6-fold respectively (Fig. 5C). In contrast, in presence of the scaffolding droplets, mean fluorescence remained and corresponding variability remained stable within 1.5-fold fluctuating range (Fig. 5C), qualitatively confirming our hypothesis. Thus, TEARS are capable of increased robustness towards both intrinsic noise and environmental changes to provide homeostasis, and reducing robustness to enhance the variability, potentially for developmental or evolutionary benefits.

### rCAG-mRNAs buffers translation efficiency against changes in RBS strength

To apply robustness tuning, we revisited the control of gene expression by fusing rCAG-repeats at 5’ UTR of mRNAs (rCAG-mRNAs) shown in Figure 1H. As we demonstrated that the output of ligand sorting is insensitive to reactant supplies, we asked whether it is also true for translational processes (Fig. 5E). We constructed CAG-RBS-gfp and RBS-gfp mRNAs using RBS strength variants (Fig. 5F). GFP expression varied across a 30-fold range in the cytosolic RBS-gfp fusions, but less than 2-fold across rCAG-RBS-gfp fusions (Fig. 5F). Thus, the output of LLPS ligand sorting is insensitive to reactant supplies and could buffer changes of ribosomal binding affinity. For biotechnological applications, rCAG-mRNAs can be used to encode proteins that require stable but not high expressions (e.g. essential genes), so that genetic drifting in RBS can be largely tolerated.

The results with wild-type RBS indicate that most rCAG-mRNA is sequestered inside the droplets (Fig. 1H), yet translations of rCAG-RBS-gfp using variants m1 - m3 are similar or even higher than respective RBS-gfp (Fig. 5F). Taken together, this suggests that translation can occur inside an rCAG-mRNA droplet, which may contribute to the majority of GFP synthesis, reminiscent of rationally-designed orthogonally-translating devices (*18*). Furthermore, once bound to the mRNA, 30S ribosomal units need to recruit other units involved in translation. Given their high turnover rates (e.g. ~ 3.7 s^−1^ for aminoacyl-tRNAs (*39*)), the ligand-sorting theory predicts highly repressed elongation inside the rCAG-mRNA droplets. Thus, initiation will not be rate limiting for translation inside rCAG-mRNA droplets, which may explain the high robustness across different RBSs.

### High-purity deoxyviolacein production by colocalization of VioABCE into TEARS

There are ample examples of *in vivo* synthetic scaffolding of metabolic pathways in recent biotechnology literature (*12, 17, 18*). Given the simplicity of TEARS engineering, and the potential advantages of two-layered structures, we focused on assembling functional TEARS made by rCAG-MS2, and used orthogonal rCAG-BoxB in control samples.

Using TEARS, we can cluster enzymes to improve product yield by accelerating the processing of unstable or toxic intermediates. Here we focused on the biosynthesis of violet-color antitumor drug deoxyviolacein by the Vio operon from *Chromobacterium violaceum* (*26*) (Fig. 6A) to benchmark our system to prior attempts. The sequential catalysis by VioABEC converts L-tryptophan into deoxyviolacein (DV) through a branching intermediate protodeoxyviolaceinic acid (PVA) (Fig. 6A, and S13A). PVA could either spontaneously turn into red-color prodeoxyviolacein (PDV), which is further dimerized into green-color deoxychromoviridans (DCV) (Fig. S13A) or transformed via a competing reaction to DV when sufficient VioC is present to catalyze this reaction. We engineered the pathway by tagging all four enzymes with tdMCP. Without clustering, overexpression of all enzyme fusions directed metabolic flux mostly toward PDV-DCV production suggesting VioC activity was not high enough to compete with the spontaneous branch; however, in presence of the cognate rCAG-MS2 for clustering all 4 VioABCE pathway enzymes, cells produced exclusively DV with undetectable (< 1%) flux towards side products (Fig. 6, B and C). In comparison, the protein-fusion based LLPS enhanced the pathway specificity 18-fold, measured by DV/PDV ratio (*17*).

**Fig. 6.**
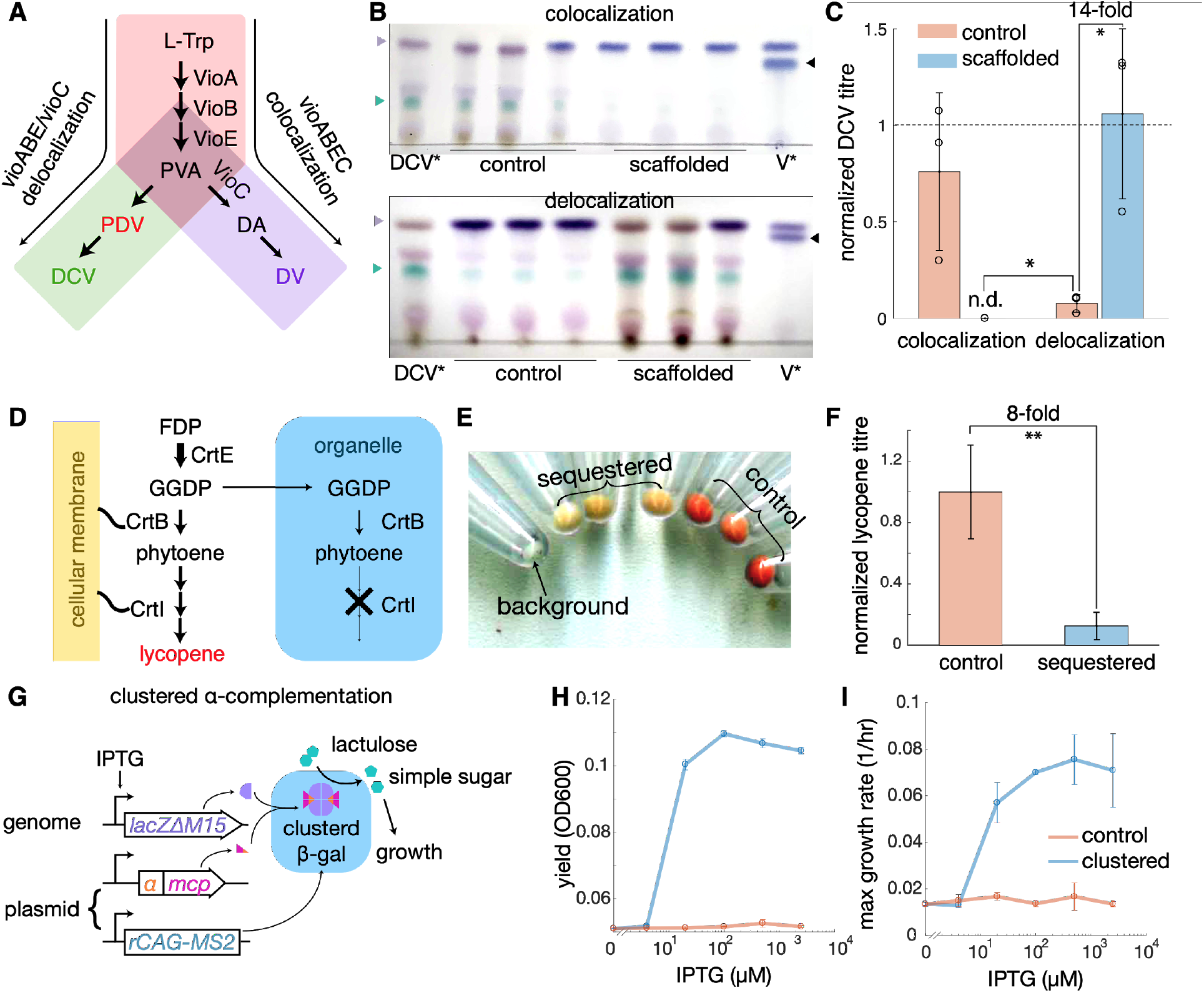
Metabolic engineering with programmed TEARS. (A and B) Colocalization and delocalization of deoxyviolacein pathways. (A) Deoxyviolacein biosynthesis has two branches: a spontaneous direction where protodeoxyviolaceinic acid (PVA) turns to (PDV, red) and then deoxychromoviridans (DCV, green), and a enzymatic branch where PVA is catalyzed into deoxyviolaceinic acid (DA) by VioC, and DA spontaneously matures into deoxyviolacein (DV). Colocalization of VioABCE (red then purple) is designed to improve deoxyviolacein biosynthesis, and delocalization of VioABE from VioC (red then green) is designed to direct metabolic into PDV-DCV direction. (B) High-performance thin-layer chromatography (HP-TLC) of extracts from strains expressing rCAG-MS2 (scaffolded) or rCAG-BoxB (control) with enzymes for colocalization (top, tdMCP-tagged VioABCE), or delocalization (bottom, tdMCP-tagged VioABE, untagged VioC). Samples from left to right are 1 μl extracts from cells only expressing rCAG-MS2 scaffolded VioABE (DCV*), 3 biological replicates each of the control and the scaffolded, and 1 mg/ml standard solution of violacein:deoxyviolacein mixture (V*). Background colors were subtracted from the image. Purple, green, and black triangles highlight the PDV/DV, DCV, and violacein. (C) Quantifications of DCV titre, normalized by extracts from DCV*, to estimate the flux towards PDV-DCV direction. N.d. not detectable. (D) Sequestration of CrtBI by TEARS to repress lycopene biosynthesis. Farnesyl diphosphate (FDP) is sequentially catalyzed into geranylgeranyl diphosphate, phytoene, and lycopene, by CrtEBI, where CrtI and potentially also CrtB are activated by membrane association. Droplets of rCAG-MS2 recruit tdMCP-tagged CrtBI thereby inhibiting their activity. (E) Pellets of cells coexpressing CrtE and tdMCP-tagged CrtBI with rCAG-MS2 (sequestered) or orthogonal rCAG-BoxB (control). 1 ml culture from 3 biological replicates for each were harvested. The background strain (DH5a) does not express the lycopene pathway. (F) Quantification of lycopene titre, normalized by the control strains. (G to I) Clustering of α-complementation. Diagram (G) of sequential recruitment of ω-fragment of β-galactosidase (β-gal) through α-complementation for sufficient catabolism of lactulose to support growth. The energy and substance requirements for growth were termed as “cost”. Yields measured by OD 600 after 50-hour culturing (H) and maximal growth rate (I) of DH5a cells expressing endogenous ω-fragment, and heterogenous α-tdMCP, with rCAG-MS2 for clustering (blue), or with rCAG-BoxB as a control (red). Errors are measured values from 3 biological replicates for all experiments. *, p < 0.05, **, p < 0.01, in paired Student’s t-test.

Thus, our modular design provides a simple approach to efficiently colocalize larger numbers of different enzymes using a single aptamer-ABD recognition pair. This presents a marked advantage over earlier attempts to assemble a long pathway using multiple orthogonal binding sites (*7*), or simple translational fusions of LLPS domains to the enzymes (*17*).

### Metabolic shunting at branch points by delocalization of VioABE and VioC

TEARS can also be programmed to sequester enzymes and shunt the metabolic flux towards the opposite direction (Fig. 6A). To demonstrate this effect, we constructed a variant containing tdMCP-tagged VioABE, and untagged of VioC that produces high amounts of deoxyviolacein. Coexpressing this system with orthogonal rCAG-BoxB produced mostly DV with small amounts of DCV; coexpressing with rCAG-MS2 completely pivoted metabolism towards DCV production as expected in a switch-like manner (Fig. 6, B and C). We confirmed that the higher flux towards DCV was not due to higher accumulation of PVA by clustering VioABE. Strains expressing VioABE with or without clustering had no significant differences in PDV/DCV production (Fig. S13B).

### Sequestration of membrane-associated enzymes to repress lycopene pathway

We next used TEARS to regulate linear metabolic pathways by delocalizing enzymes away from their subcellular localization, focusing on biosynthesis of yellow-to-red-color antioxidant lycopene production by CrtEBI from *Erwinia uredovora* (Fig. 6D). It was previously shown that *E. uredovora* CrtI is only active when binding to phospholipid membranes (*27*). Although CrtB was not well-characterized in bacteria, its homologous phytoene synthase in plants also requires membrane association to be functional (*40*). We engineered this pathway by tagging CrtBI with tdMCP and kept CrtE untagged. Coexpressing engineered enzymes with orthogonal droplets (rCAG-BoxB) results in visibly red-color bacteria (Fig. 6E). Despite the potential competition between membranes and droplets to bind CrtBI, sequestering CrtBI away from CrtE by the cognate rCAG-MS2 droplets significantly reduced lycopene titre by 8-fold on average (Fig. 6F, and S14), and resulted in yellow-color bacteria (Fig. 6E). Therefore, TEARS can be used to repress metabolism by a spatial “knockout” of essential enzymes in the pathways.

### Clustered α-complementation rescues growth defect of lacZΔM15 mutant strains

The ligand-sorting framework suggested a method to organize endogenous components by synthetic receptors with low turnover and/or high binding rates. As a proof of concept, we designed a system to rescue bacterial growth of *lacZ*ΔM15 mutants on an artificial sugar lactulose (*41*). Lactulose is imported by *lacY*-encoded lactose permease, and then catabolized by *lacZ*-encoded β-galactosidase (β-gal), but unlike lactose, it does not induce expression of the lac operon. In *lacZ*ΔM15 mutants, the genome only encodes the ω-fragment of LacZ with no activity. Additional expression of the *lacZ*α-encoded α-peptide renders β-gal with partial activity (*28*). We expressed the α-peptide-tdMCP fusion (α-tdMCP), in presence of the cognate rCAG-MS2 droplets to sequentially recruit ω-fragments, catalyze the α-complementation, and concentrate functional β-gal (Fig. 6G, and S15, A-D).

Overexpressing α-tdMCP from a high-copy plasmid is not sufficient to support the strain’s growth on lactulose, even under full induction of genomic *lacZ*ΔM15 (Fig. S15E). In contrast, scaffolding of α-tdMCP using the cognate rCAG-MS2 droplets enabled growth under IPTG induction ≥ 20 μM; as control, replacing the rCAG-MS2 with the orthogonal rCAG-BoxB droplets had no rescue effects (Fig. 6, H and I). Thus α-complementation of β-gal can rescue growth defects of *lacZ*ΔM15 mutants under lactulose regime only when it was organized by droplets to enhance the complementation, and to concentrate the complemented enzymes.

## Discussion

Our work establishes a robust approach to harness functional synthetic RNA-based condensates for fundamental understanding of LLPS in as much as for metabolic engineering. The modular design of TEARS significantly simplifies the design-build-test cycles so that each part of the droplets can be engineered independently. We proved that rCAG-repeat RNAs are sufficient for LLPS so that any desired solutes can be enriched by aptamer-ABD binding. The system design can be further complexified by mixing different rCAG aptamers or by recruiting oligomerizing proteins (*12*). It will therefore be an interesting subject for future characterization to create a more generic design principle.

Using TEARS as a model of LLPS, we observed behaviors that were not previously characterized with in vitro experiments, or with systems only containing a single LLPS protein. These include simple mechanisms for multiphase separation and compositional control, and diverse regulations of concentration robustness. Our results may provide further insight to the origins and functionality of natural condensates. For instance, though often overlooked, fluorescent protein labelling of eukaryotic condensates often results in foci void of fluorescence (*12, 16, 33*), which is reminiscent of our observations of multilayering. The compositional control by ligand sorting may be relevant to natural organelles, such as ribonucleoprotein granules (*20*) that rapidly shift their preferences in response to growth conditions. Turnover-governed preferences over the same pair of interacting molecules may explain how systems like nucleoli (*16, 33*) select reactants over structurally-alike products. Moreover, we demonstrate that condensates may have diverse effects on cellular phenotypic variability beyond the established noise reduction role. These include the possibility and critical conditions to amplify noise, but also to provide robustness against large changes in solute concentrations or protein expression parameters suggested by the theoretical analysis.

Our main incentive to build LLPS in bacteria was to engineer intracellular reactions. The remarkable efficiency of TEARS to exclude cytosolic components provides an unprecedented environment amenable for application purposes. We showed simple methods to colocalize a long pathway for metabolic channeling of deoxyviolacein production, which resulted in optimal desired efficiency at the branch point. We also demonstrated delocalization for metabolic shunting or enzyme sequestration. With clustering of α-complementation, we illustrated enhancement of endogenous proteins through sequential recruitments. Particularly, all of the systems used the same rCAG-MS2 construct, and none of them required fine-tuning of expression levels, proving the reusability of our design. We anticipate this would inspire further applications for more complex organization and repurposing for other chassis beyond *E. coli*.

## Acknowledgments

We are grateful for the support received from C. Lotton and other INSERM U1001 and Center for Research and Interdisciplinarity members. We specifically thank Dr. J. Wintermute for critical reading of the manuscript and for the TEARS acronym. We thank members from the Paris Bettencourt iGEM 2017 team for their support and comments for the early development of the project. We thank A. Jain for sharing pHR-Tre3G-47xCAG-12xMS2 plasmid. We thank the members of INSERM U1001, Center for Research and Interdisciplinarity, and D. Bikard, Y. Ponty for critical comments on earlier versions of the project, R. Kusters and M. Chen for comments on modeling, and C. Li and E. H. Wintermute for the comments on the manuscript.

## Funding

This project has received funding from the European Union’s Horizon 2020 Research and Innovation Programme under the Marie Sklodowska-Curie Grant Agreement No. 665850 and was further supported by the Bettencourt Schueller Foundation.

## Author contributions

H.G conceived the study and experiments. H.G. and J.C.R. performed molecular cloning, and optical imaging. J.C.R. and A.M. perform TEM imaging. H.G. and V.P. performed purification assay. H.G. performed flow cytometry and growth measurement, and experiments for metabolic engineering. H.G and A.D cloned the ELK16 system. H.G. and X. Song performed imaging analysis. H.G. developed the theoretical analyses, and performed the numerical analysis. A.B.L mentored INSERM U1001/U1284 project members. H.G and A.B.L interpreted data and wrote the manuscript with inputs from J.C.R, X. Song, and A.M.

## Supplementary Materials

Materials and Methods

Supplementary Text

Figures S1-S15

Tables S1-S3

References 42-45

## Notes

### Competing Interest Statement

The authors have declared no competing interest.

### Summary of Updates

Remove the acronym from the title

