## Supplementary material for "Spatial engineering of *E. coli* with addressable phase-separated RNAs": Materials and Methods, Supplemental Text, Figures and Tables

**This PDF file includes:**

Materials and Methods
Supplementary Text
Figs. S1 to S15
Tables S1 to S3

### Materials and Methods

Plasmid and cloning methods

Standardized parts (named after "BBa_") and backbones (named after "pSB") documented in iGEM part registry (https://parts.igem.org/) were used in this study. Customized genetic parts (listed in Table S1) were derived from PCR of the given templates, annealed from oligos (Integrated DNA Technologies), or de novo synthesized as gblocks (Integrated DNA Technologies) and gene fragments (Twist Bioscience). Designated backbones are listed Table S2. Customized backbone were modified from an original backbone, by restriction-enzyme-based cloning or Gibson assembly (New England Biolabs). Inserts < 300 bp were cloned by oligo annealing. Inserts between 300 bp and 3kb were cloned as one piece of fragment into designated backbone using Gibson assembly or Golden Gate assembly through BsaI-HFv2 (New England Biolabs). Inserts > 3kb were first sub-cloned into high-copy backbone pSB1C3 or pSB1A3, and then assembled into designated plasmids using Gibson assembly or Golden Gate assembly. Sequences of inserts were verified by Sanger sequencing, and all bases were covered in at least one reaction. All the final plasmids characterized in experiments with sequences and notations were listed in Table S3.

Strains, growth medium, and induction conditions

*E. coli* strain "NEB stable" (New England Biolabs) was used to clone rCAG-BoxB. *E. coli* strain DH5a (taxid: 668369) was used for all the routine cloning. *E. coli* strain BL21AI (Invitrogen) was used for imaging unless another strain was indicated. BL21AI was also used for characterization of Vio pathway assay. DH5a was used in cytometry assays, characterization of lycopene assay, and clustering α-complementation assay. E. coli strain MG1655 was used as the positive control in clustering α-complementation assay. To transform over two plasmids, two plasmids were co-transformed first to create an intermediate strain, and then the rest of the plasmids were transformed into the intermediate strain. To avoid deletions due to repetitive sequences, or mutations due to toxicity of metabolic pathways, we re-sequenced plasmids after transformation. We occasionally observed further extensions of the CAG repeats; these did not significantly change the phenotypes. We did not observe any large deletions as previously reported when cloned into high-copy plasmids (*22*).

LB-Miller medium (BD Difco) was used for strain propagation, bacteria culture to mid-exponential phase, and culture for imaging, flow cytometry analysis, mother machine, Vio pathway production, and lycopene production. M9-lactulose minimal medium (1× M9 salts (BD), 2 mM MgSO4, 0.1 mM CaCl2, 0.4% lactulose (Sigma-Aldrich)) was used in the α-complementation clustering assay. 100 μg/ml ampicillin, 50 μg/ml kanamycin, and 30 μg/ml chloramphenicol were used for antibiotic selections. Unless specified, 100 ng/ml anhydrotetracycline (aTc), 0.1 mM IPTG, and 0.2% L-arabinose were used to induce genes under tetracycline-inducible promoters, LacI-repressed promoters, and AraC-repressed promoter, respectively.

Pre-culture for imaging experiments

For imaging experiments, bacteria were first prepared through the following process to homogenize the bacterial physiology. Transformations were plated, or glycerol stocks were streaked, on plates containing LB and 2% agar (BD Difco) and relevant antibiotics but without any inducers, and grown overnight at 37°C. Plates were stored at 4°C for no more than a week. Single colonies were inoculated in 5 ml LB with a proper combination of antibiotics, and incubated at 37°C 150 rpm overnight. The overnight cultures were then diluted 200 or 250-fold in 5 ml LB with the same antibiotics to initial OD600 of about 0.05. Diluted cultures were grown back to the mid-exponential phase (OD600 = 0.3 ~ 0.5) at an indicated temperature, 150 rpm. The resulting cultures are called pre-culture at the given temperature.

Transmission electron microscopic (TEM) imaging

For transmission electron microscopy, indicated strains were pre-cultured at 30°C. Then, pre-cultures were directly supplied with aTc, and induced overnight. Overnight bacteria were fixed with 2.5% glutaraldehyde in 1M HEPES buffer pH 7.2 overnight at 4°C. Specimens were postfixed 1h in 1% osmium tetroxide in 1M HEPES buffer pH7.2 and were dehydrated in a graded series of ethanol, and embedded in Epon. After polymerization, thin sections were cut with an Ultramicrotome (Leica, Ultracut UC7). Sections were stained with uranyl acetate at 4% and lead citrate. Images were acquired with a TEM Tecnai SPIRIT at 120 kV accelerating voltage with a camera EAGLE 4K x 4K (FEI-Thermofisher Company).

Standard phase-contrast and fluorescent microscopic imaging

For imaging of bacteria induced overnight (Fig. 1D, and 4E), samples were prepared in the same process as for TEM imaging. For imaging of cells induced at exponential phase (Fig. 1 to 3, and 4D), bacteria were first pre-cultured at indicated temperature (for ABD-GFP imaging, 37°C by default, or 30°C if noted; for BiFC systems, 30°C). Then, bacteria were induced by directly adding aTc, for another 2 hours at 150 rpm in the same temperature. To be noticed, we did not induce the production of PN-GFP by IPTG. Leaky expression was sufficient for imaging. For imaging of BiFC systems in log-phase cells under different inductions (Fig. 5E), bacteria were first pre-cultured at 30°C. Then, pre-cultures were adjusted OD600 = 0.3 in 200 µl LB supplied with antibiotics, 0.1 mM IPTG, 50 ng/ml aTc and arabinose from 0% to 0.2% (w/v) by 5-fold serial dilution, in standard U-bottom 96-well plates (Greiner Bio-one), sealed by aluminum foil (Thermo Fisher), and incubated at 30 °C 750 rpm in Titramax 1000 (Heidolph) for 2 hours. 1 µl of samples were placed between a cover slip and 1% agarose pad. Images were acquired using a Nikon Ti-Eclipse microscope, CFI Plan Apo Lambda 100X Oil objective (NA=1.45) and ProEM+:1024B EMCCD camera, controlled by Metamorph software, in phase contrast, and fluorescence channels (GFP excitation 480/30, dichroic mirror 505, emission 535/45; mKate excitation 560/40, dichroic mirror 600, emission 635/60). Focus was readjusted manually for every position.

Pulse-chase analysis by time-lapse imaging in mother-machine microfluidic

The mother machine microfluidic setup used was detailed in our previous study (*31*). Strain pre-cultured at 37°C were loaded into the channels. LB supplied with chloramphenicol, kanamycin, and aTc were constantly flowed into the microfluidic devices using Hamilton GC-grade glass/PTFE syringes (Gastight 1000 Series) and a high-precision syringe pump (Harvard Apparatus PHD 2000 Programmable). Microfluidic chips were incubated at 37°C, and analyzed using previously described microscopy setup. Focus was readjusted by Nikon's perfect focus system before each imaging cycle for every position. After visible fluorescent foci were formed, phase-contrast and GFP images were acquired for every position once every 10 minutes for up to 10 hour. Due to cell adaptation mediated by the acrAB-TolC (*32*), cells were first induced and formed the droplets, and then rapidly adapted to the aTc level and lost expression.

Segmentation analysis of phase contrast and fluorescent images and statistical analysis

Experiments were repeated by H.G and J.C.R independently in different days to verify reproducibility. Images from the same batch were randomly chosen for analysis by X.S. Sample sizes were not predetermined by any statistical methods. During the imaging analysis and statistical analysis, the investigator was blinded to the allocation. Images were processed using a standardized pipeline (*42*). Data acquired from segmentation was further analyzed using Matlab. For cells containing TEAR droplets, the "cytosol" is considered as the area within the cell excluding all identified foci. To be noticed, automatic analysis may include solute-free phases in TEAR droplets into the cytosol area as well. For all analyses, cells, foci, or cytosol that have negative mean fluorescence intensity were discarded from the dataset. We denote the fluorescence values of a subject of interest (e.g. cytosolic fluorescence), as value $F$. For mean fluorescence estimation, we bootstrapped data to calculated mean fluorescence $\langle F \rangle_i$, $i$ from 1 to 10,000. For bootstrapping, we first calculated the mean and standard deviation of fluorescence on logarithm scale as $\langle \log(F) \rangle$ and $\sigma(\log(F))$, discard the outliers that 2.5 standard deviation away, i.e. $|F - \langle \log(F) \rangle| > 2.5\ \sigma(\log(F))$, and then bootstrapped data to calculate coefficient of variation $CV_i(F) = \sigma(F)/\langle F \rangle$. Then we calculated expected values and standard deviation of bootstrapped results. To be noticed, keeping outliers during bootstrap can cause inappropriate distributions.

Condensate purification assay

To verify reproducibility, the following protocol was repeated by H.G and V.P independently, over 3 times using the BL21AI strains, and once with DH5a strains for each condition. Indicated strains were grown overnight 37°C. Overnight culture were diluted in 5 ml LB and antibiotics, to initial OD600 ~ 0.005, and then incubated at 37°C 150 rpm to OD600 ~ 0.3 - 0.4. Cultures were supplied aTc to final concentration 100 ng/ml, and then incubated for another 5 hours. The induced cultures were centrifuged 3,000 rpm for 10 mins. Pellets were resuspended in 1 ml PBS, and centrifuged again 6000 rpm 4 mins. Supernatants were discarded and remained pellets can be lysed immediately or stored in -80°C for future use. To lyse the cells, pellets were completely resuspended in 5 ml per gram of wet cell pellet of detergent BugBuster® Protein Extraction Reagent (Merck Millipore), at room temperature by repetitive pipetting. Lysates were centrifuged, and the supernatants were isolated and denoted as "supernatant" fraction (Fig. S1C). Then the remaining pellets were processed with lysozyme (Sigma-Aldrich), and centrifuged again, of which supernatants and pellets were separated and denoted as "cell debris" and "condensate", respectively (Fig. S1C). Condensates were washed twice by 1:10 diluted BugBuster reagent to remove remaining components from cell debris. 5 µl of supernatants, cell debris, wash, purified condensates from indicated strains are denatured by adding 5 µl 2 × Laemmli Sample Buffer (Bio-Rad) at 95°C for 10 mins. 5 µl of denatured samples, together with 5 µl Precision Plus Protein™ Dual Color Standards (Bio-Rad), are loaded in Mini-PROTEAN Precast SDS-PAGE Gels (Bio-Rad). SDS-PAGE gels are stained by GelCode Blue Safe Protein Stain (Thermo Fisher Scientific).

Flow cytometry assay

Three biological replicates for indicated strains were inoculated into 0.75 ml LB with chloramphenicol, in 2-ml 96-well deep-well plates (Thermo Fisher Scientific). The plates were sealed by breathable films (Greiner), and incubated in 37°C 750 rpm overnight in a Multitron shaker (INFORS HT). Overnight bacteria were diluted 1:250 in 750 µl fresh media in 2-ml deep-well 96-well plates, and grown at 37°C 750 rpm for 3 hours. 2 µl cultures of log-phase cells were diluted in 750 µl LB with chloramphenicol, and 100 ng/ml aTc. After mixing by pipetting, 100 µl cultures of each were transferred into a standard 96-well plate, sealed by breathable film, and incubated at 37°C 750 rpm overnight. 10 µl overnight cultures were in 100 µl PBS in standard 96-well plate, and measured by BD LSR Fortessa™ X-20 cell analyzer with High Throughput Sampler under high-throughput screening mode. At least 1000 events were acquired for each sample. Medians were used as the representative statistic of fluorescence intensity for all the samples. Background auto-fluorescence was not subtracted. Relative fluorescence units (RFU) were calculated by the ratio of fluorescence intensity of sample of interest over the auto-fluorescence intensity.

Growth curve measurement of strains expressing rCAG-MS2 or tandem MS2 repeats alone

Three biological replicates of two strains were pre-cultured at 30°C, in the same process used for imaging. For growth curve measurements, pre-cultures were adjusted to the same OD, and then diluted 1:100 in LB medium supplied with antibiotics, and indicated ng/ml aTc. 100 µl cultures of each were transferred into 96-well µClear black plates. Plates were incubated in 30°C, continuous, linear shaking (amplitude 3 mm, frequency 540 rpm) in Tecan SPARK plate reader. OD at 600 nm were measured every 10 mins overnight. The OD values between two strains were fitted by the identify line, and R-square values were calculated accordingly (Fig. S12), using Matlab.

Vio pathway assay

Three (Fig. 6B) or five (Fig. S10B) biological replicates of indicated strains were cultured overnight at 37°C 150 rpm in 5 ml LB with antibiotics. Then 25 µl overnight cultures of each replicate were diluted in 5 ml LB supplied with antibiotics and 100 ng/ml aTc, and incubated at 37°C 150 rpm for 24 hours. 24-hour cultures were centrifuged 4500 rpm for 10 mins, resuspended by 1 ml PBS, and then centrifuged 15000 rpm for 10 mins.

For thin layer chromatography (HP-TLC), pigments in pellets were extracted by mixing with 500 µl methanol for ~ 1 hour. 1 µl of each sample was spotted on 50×75 mm unmodified silica gel 60 TLC plates (VMR). As positive controls, same volumes of extraction of strains only expressing tdMCP-tagged VioABE, i.e. pVioAMBMEM (DCV*), and a 1 mg/ml standard solution of violacein:deoxyviolacein mixture (85:15, w/w) (Sigma-Aldrich) (V*) in 1 ml methanol were sampled on the two sides of the silica gel. Plates ran vertically for ~ 3 mins in chloroform:methanol (6:1, v/v). Silica gels were imaged under indoor light, and quantified by Matlab. Production of DCV was estimated by the area of its peak.

For plate reader measurements of PDV/DCV production, pigments in pellets were extracted by 200 µl DMSO, vortexed, and then centrifuged again. 100 µl DMSO supernatants were transferred into 96-well µClear black plates (Greiner), and OD from 300 to 800 nm were measured by Tecan SPARK plate reader. OD at 588 nm were used to represent the amounts of PDV/DCV production. Curve profiles of DCV* and V* were used to qualitatively estimate the type of dominant compound in the extract mixtures.

Isolation of VioDΔ(264-372)

For the delocalization assay, a control strain was required to have high efficiency to produce deoxyviolacein for proper comparison. We chose not to change promoters, RBS, or coding sequences of VioABCE, so that concentrations and activities of enzymes in colocalization and delocalization assays are comparable. Considering that expressions of VioABCDE produce violacein and deoxyviolacein with low amounts of DCV production (*26*), we originally chose to construct plasmids constitutively expressing tdMCP-VioD in addition to untagged VioC to demonstrate delocalization. Cassettes of untagged VioC and tdMCP-tagged VioD, both translated by BBa_B0034 in a bicistronic cassette driven by BBa_J23105, were cloned in pSB1A3. This library was transformed into DH5a. Random clones were inoculated, and used to purify individual plasmids. Considering the toxicity of violacein (*26*), we tested whether the plasmids can be stably propagated one-by-one by co-transforming them with pVioAMBMEM and the orthogonal RNA rCAG-BoxB (Table S3). Violet-color (as judged by eyes) colonies were inoculated and analyzed by plate readers (see Vio pathway assay). Plasmids resulting in high violet pigment production were then analyzed by Sanger sequencing. One plasmid pVioCD53 tested in Variant No. 53 showed C-terminal truncation from 264 aa (Table S3), probably due to intra-plasmid recombination. Further HP-TLC analysis showed that only deoxyviolacein but no violacein were produced in Variant No. 53. It suggests that truncated VioD, although lost catalytic activity, may serve as an enhancer for VioC. We used pVioCD53 for delocalization assay instead of original design, so that the final product is still deoxyviolacein.

Lycopene pathway assay

Three biological replicates of indicated strains were inoculated in 500 μl LB, with ampicillin, chloramphenicol, in 2-ml deep-well 96-well plate, sealed by breathable film, incubated 37°C , 750 r.p.m. in multitron overnight. 60 μl overnight cultures were diluted in 3 ml fresh media, and grown in 28°C 150 rpm to mid-exponential phase. 1 ml log-phase culture was diluted in 50 ml LB supplied with ampicillin, chloramphenicol, and 100 ng/ml aTc, in 250 ml flask, and incubated in 28°C 150 rpm for 48 hours. For imaging of cell pellets, 1 ml 48-hr culture were centrifuged 10,000 rpm for 1 mins, and supernatants were discarded. For quantification by plate reader measurements, 1 ml 48-hr cultures were centrifuged 15,000 rpm for 1 minute. Pellets were resuspended with 200 μl DMSO, mixed in room temperature for 1 hour, and then centrifuged at 15000 rpm for 10 minutes. 100 μl DMSO supernatants were transferred into 96-well μClear black plates, and OD was measured from 300 nm to 800 nm with a Tecan SPARK plate reader. As a positive control, a standard solution of lycopene (Sigma-Aldrich) in DMSO was measured to calibrate the reading. The ratio of an indicated sample over OL1 was estimated by calculating the ratios of OD values from 400 nm to 550 nm, and then calculating the mean of ratios.

Clustering α-complementation assay

To measure growth curves, three biological replicates of indicated strains were pre-cultured to exponential phase at 37°C, in the same process used for imaging. Pre-cultures were adjusted to the same OD, and then diluted 1:100 in M9-lactulose minimal medium supplied with antibiotics, 25 ng/ml aTc, and indicated IPTG. To be noticed, the final media is actually LB: M9-lactulose 1:100. 100 μl cultures of each were transferred into 96-well μClear black plates. Plates were incubated in 37°C, continuous, linear shaking (amplitude 3 mm, frequency 540 rpm) in Tecan SPARK plate reader. OD at 600 nm were measured every 10 mins for 350 cycles. Transient growth rates were defined as the local derivative of the logarithm OD600, and calculated by linear regression of every 11 data points. Maximal growth rates were therefore calculated as the maximum of transient growth rates of the given curves. Yields were calculated as the increase of OD after 50 hours.

Watercolor illustration

Biomolecules illustrated by watercolor in schematic diagrams (Fig. 1A, 1B, and 4A) were produced by Illustrate (*43*). Templates were acquired from RCSB Protein Data Bank (PDB):

single-stranded and duplex CAG repeats, 5VH7; MS2 aptamer, 1ZDH; tdMCP, 1ZDH; BoxB aptamer, 1QFQ; full-length Protein N, 6GOV; RBS, 1SDR; 30S ribosomal unit, 1ML5; FA, FB, and complemented GFP, 1HUY.

**Supplementary Text**

1. Minimal model of well-mixed and organelle-organized reactions in a mesoscopic scale

1.1. Mass-action description of well-mixed reactions

Here we use the simplest two-species reaction as a representative example and solve their kinetics according to the law of mass action. The results we acquired from this model will be used as the benchmark when we compare with the system organized by organelles (membrane-enclosed, or phase-separated). Chemical species under considerations are:

(1) R, receptor;
(2) L, ligand;
(3) RL, receptor-ligand complex;
(4) P, product.

We first consider a case that ligand-receptor reaction is irreversible. The chemical reactions in the system includes:

$$0 \underset{\alpha_R[R]}{\overset{\beta_R}{\rightleftharpoons}} \mathrm{R} \tag{R1}$$

$$0 \underset{\alpha_L[L]}{\overset{\beta_L}{\rightleftharpoons}} \mathrm{L} \tag{R2}$$

$$\mathrm{R} + \mathrm{L} \underset{k'[RL]}{\overset{k[R][L]}{\rightleftharpoons}} \mathrm{RL} \xrightarrow{k_c[RL]} \mathrm{P} \tag{R3}$$

$$\mathrm{RL} \xrightarrow{\alpha_{RL}[RL]} 0, \mathrm{P} \xrightarrow{\alpha_P[P]} 0 \tag{R4}$$

where "0" represents the systems that are not included; [X] is concentration of the species X, with a unit as molecules per unit volume; $\beta_X$ and $\alpha_X$ are rate constants for synthesis and turnover (dilution, degradation, or consumption by other reactions) of species X; $k$ and $k'$ are the rate constant for reversible association and dissociation between ligand and receptors. $k_c$ is the irreversible rate constant of the maturation from ligand-receptor complexes to the final product. A system containing only reversible reactions, can be considered as a special case for irreversible systems where the final step of maturation in R3 does not exist.

1.1.1. Dynamics of well-mixed reaction and steady-state solutions

Here we described the well-mixed system by a set of ordinary-differential equations (ODE):

$$\begin{aligned}
\frac{d}{dt}[R] &= \beta_R - \alpha_R[R] - k[R][L] + k'[RL] \\
\frac{d}{dt}[L] &= \beta_L - \alpha_L[L] - k[R][L] + k'[RL] \\
\frac{d}{dt}[RL] &= k[R][L] - k'[RL] - \left(k_c + \alpha_{RL}\right)[RL] \\
\frac{d}{dt}[P] &= k_c[RL] - \alpha_P[P]
\end{aligned} \quad . \tag{1}$$

By defining an “apparent” rate constant to describe the net association of ligands and receptors, as the following:

$$k_i = \frac{k\left(k_c + \alpha_{RL}\right)}{k' + k_c + \alpha_{RL}}$$

the steady-state solution of these ODEs can be given by:

$$[R]^* = \frac{1}{2\alpha_R}\left(\sqrt{\Delta} - \frac{\alpha_R\alpha_L}{k_i} + \left(\beta_R - \beta_L\right)\right)$$

$$[L]^* = \frac{1}{2\alpha_L}\left(\sqrt{\Delta} - \frac{\alpha_R\alpha_L}{k_i} - \left(\beta_R - \beta_L\right)\right)$$

$$[RL]^* = \frac{1}{k_c + \alpha_{RL}}\left(\frac{1}{2}\left(\frac{\alpha_R\alpha_L}{k_i} + \beta_R + \beta_L\right) - \frac{1}{2}\sqrt{\Delta}\right)$$

$$[P]^* = \frac{1}{\alpha_P}\frac{k_c}{k_c + \alpha_{RL}}\left(\frac{1}{2}\left(\frac{\alpha_R\alpha_L}{k_i} + \beta_R + \beta_L\right) - \frac{1}{2}\sqrt{\Delta}\right) \tag{2}$$

where $(.)^*$ represents the steady-state values, and:

$$\Delta = \left(\frac{\alpha_R\alpha_L}{k_i} + \beta_R + \beta_L\right)^2 - 4\beta_R\beta_L \,. \tag{3}$$

Here $\Delta$ is the discriminant of the system, which is always > 0 by definition.

1.1.2. Approximation under simplified conditions.

We assume all solutes are stable against degradation. Turnover of solutes are mainly governed by dilution caused by exponential growth, i.e. $\alpha_R = \alpha_L = \alpha_{RL} = \alpha_P = \alpha$. And the synthesis rates are equal for receptors and ligands as $\beta_R = \beta_L = \beta/2$. Then the discriminant of the system is given by:

$$\Delta = \left(\frac{\alpha^2}{k_i} + \beta\right)^2 - \beta^2 \,. \tag{5}$$

Given the symmetry of the system, receptors and ligands have the same concentrations. Thus, the apparent binding rate of ligand and receptors in a mixed reactions ($v_M$) at the steady state is given by:

$$v_M = k_i[R][L] = \frac{1}{2}\left(\frac{\alpha^2}{k_i} + \beta\right) - \frac{1}{2}\sqrt{\left(\frac{\alpha^2}{k_i} + \beta\right)^2 - \beta^2} \,. \tag{7}$$

1.2. Mesoscopic reaction-diffusion description of irreversible ligand sorting by organelles

Now we consider a system that separates into three phases, an aqueous phase (a.q. for short, e.g. cytosol, nucleosol) with a volume fraction as $\phi_A$, and an organelle including the solute-rich phase with a small volume fraction as $\phi_R \ll 1$, and a solute-free phase $\phi_P$. By definition, $\phi_A + \phi_R + \phi_P = 1$.

For the simplicity of the model, we assumed the concentration of RNA solvents is relatively low in aqueous phase, thus ignoring the effects of receptor-solvent association in the following deduction. Therefore, the same four chemical elements are under consideration, receptors, ligands, receptor-ligand complex, and final products.

For each species X, two states of species exist: one in the aqueous phase, denoted as $X_A$, and one in the solute-rich phase of the organelle, denoted as $X_O$. We consider reactions in the aqueous and organelles phases still follows mass-action laws as previously described:

$$0 \underset{\alpha_R[R_A]}{\overset{\beta_R}{\rightleftharpoons}} R_A \tag{R5}$$

$$0 \underset{\alpha_L[L_A]}{\overset{\beta_L}{\rightleftharpoons}} L_A \tag{R6}$$

$$R_A + L_A \underset{k'[RL_A]}{\overset{k[R_A][L_A]}{\rightleftharpoons}} RL_A \xrightarrow{k_c[RL_A]} P_A \tag{R7}$$

$$R_O + L_O \underset{k'[RL_O]}{\overset{k[R_O][L_O]}{\rightleftharpoons}} RL_O \xrightarrow{k_c[RL_O]} P_O \tag{R8}$$

$$RL_A \xrightarrow{\alpha_{RL}[RL_O]} 0,\ P_A \xrightarrow{\alpha_P[P_O]} 0,\ R_O \xrightarrow{\delta[R_O]} 0,\ L_O \xrightarrow{\delta[L_O]} 0,\ RL_O \xrightarrow{\delta[RL_O]} 0,\ P_O \xrightarrow{\delta[P_O]} 0 \tag{R9}$$

where $\delta$ is the dilution rate constant of organelles, considering that organelles efficiently exclude the endogenous proteins (Fig. 1F), and thus have relatively low degradation and undesigned consumptions of the species.

Between aqueous and organelles phases, the molecule exchanges are not described by mass action, but the influx and efflux on the surface:

$$R_A \overset{J_R}{\rightleftharpoons} R_O$$

$$P_A \overset{J_P}{\rightleftharpoons} P_O \tag{R10}$$

$$R_O + L_A \overset{J_L}{\rightleftharpoons} RL_O \xrightarrow{k_c[RL_O]} P_O \tag{R11}$$

where $J_X$ is the net intake current of the species X into the organelle through the absorption of the surface organelles, in molecules per unit time.

1.2.1. Exchanges of molecules between aqueous phase and organelles

On the surface of the organelle, there is an exchange of solvent, solvent-bound receptor or product, through solvent-solvent interactions. Solvent-free receptors or products can enter the organelle through the free binding sites of the organelles, of a partition coefficient $\varepsilon$. Ligands can enter the organelle through ligand-receptor interactions. Here we first consider the intake current of ligands on the surface of the organelles.

Given the liquid nature, organelles have non-compressible volumes. Thus we define the concentration of recruitment domains (aptamers for instance) inside organelles, with unit as moles per unit volume, as a constant: $1/V$, where $V$ is the average volume of organelles divided by the number of recruitment domains they contain. We also define the average radius of solvents per each recruitment domain as $r$, that of receptors as $r_R$, that of ligands as $r_L$, and that of products approximately as $(r_R + r_L)$.

Using electrostatic analogue (*44*), we define the maximal "capacitance", $a$, of an isolated organelle of that size and shape as a measurement of the sum of its maximal capability to absorb any molecules that reaches the surface of organelle. For a spherical droplet, or more commonly in our experiments, a meniscus surface, the capacitance $a$ is simply the radius, ~ 400 nm (Fig. S4). Now considering that a certain species y enter the organelle when they hit and associate with the cognate component x on the surface of organelle, while the reverse reaction occurs at the same time that xy complex dissociate and release molecule y back to the aqueous phase:

$$\mathrm{y_A} + \mathrm{x_O} \overset{J_y}{\rightleftharpoons} \mathrm{xy_O}.$$

Then derived from Berg-Purcell model (*44*), the intake current $J_y$ of a certain molecule y through the component x on the surface of organelle, in molecules per unit time, can be given by:

$$J_y = 4\pi D_y \left( \eta_{xy} [y_A] \frac{N_x r_x a}{N_x r_x + \pi a} - \eta'_{xy} \frac{N_{xy} r_{xy} a}{N_x r_x + \pi a} \right) \tag{8}$$

where:

$D_y$ is the diffusion coefficient of species y in the aqueous phase.

$N_x$ and $N_{xy}$ are the number of free x and xy complexes on the surface of organelle.

$r_x$ is the radius of the y-binding domain in x, and $r_{xy}$ is the radius of x-y-interaction domains in x-y complexes.

$\eta_{xy}$ is the ratio between the number of x-y binding events over the number of x-y collisions, and can be defined by x-y binding rate constant as:

$$\eta_{xy} = \frac{k_{xy}}{4\pi \left( D_x + D_y \right) \left( r_x + r_y \right)},$$

so that the first term in the equation describes the current of influx considering that different receptors will compete with each other; and $\eta'_{xy}$ is the ratio between the number of x-y complexes breakdown events per unit time, over the number of ligand that can diffuse unit volume per unit time, and can be defined by x-y dissociation rate constant as:

$$\eta'_{xy} = \frac{k'_{xy}}{4\pi \left( D_x + D_y \right) \left( r_x + r_y \right)},$$

so that the second term in the equation describes the current of efflux considering that the dissociated ligands can be reabsorbed by the free receptors around it. $N_x r_x + \pi a$ measures the competition between x on the surface of organelles to absorb y from the aqueous phase, and also how many y released from organelles will be absorbed again before they are considered as in the aqueous phase.

If two molecules x, and y have similar sizes and shapes, then we assume $D_x = D_y$ , and $r_x = r_y$, and:

$$\eta_{xy} = \frac{k_{xy}}{16\pi D_y r_x} \tag{9}$$

In addition, by definition, the number of recruitment domains on the organelle surface is $N = a^2/r^2$, and that of a species i is $N_i / N = [i_O] / (1/N_A V) = [i_O] N_A V$. Thus the intake current can be rewritten into:

$$J_y = \frac{a}{4r_x} \left( \frac{k_{xy} [y_A][x_O]}{[x_O] + K_y} - \frac{k'_{xy} [xy_O]}{[x_O] + K_y} \right) \tag{10}$$

where the affinity of organelles to species y is defined as:

$$K_y := \frac{\pi r^2}{V r_x a}. \tag{11}$$

Particularly for ligand intake, the net current is given by:

$$J_L = \frac{a}{4r_R}\left(k[L_A]\frac{[R_O]}{[R_O]+K_L} - k'\frac{[RL_O]}{K_L}\right); K_L = \frac{\pi r^2}{V r_R a}. \tag{12}$$

1.2.2. Partition of receptors and products with dominant binding kinetics

Next, we consider an extreme case of molecule exchanging, where the “primary” solutes are recruited through high binding affinity and high binding rate constants. If the primary solute is not interacting with another primary solute, then we assume that they can reach detailed balance in the partitioning between organelle and aqueous phases even before other reactions in the system occur. Despite the existence of other reactions, the partition of primary solutes can always be considered as in equilibrium. This assumption, we referred to as “quasi-thermostatic partitioning”. Thus, the partition coefficient $\varepsilon$ is defined as the ratio of concentrations of primary solutes between organelle and aqueous phases:

$$\varepsilon = \frac{[x_O]}{[x_A]}. \tag{13}$$

Given proper flexible linkers between domains of those solutes, solutes that contain the same binding domain to the recruitment domains on the solvent, should be always partitioned between organelle and aqueous phases in the same way. Particularly in the ligand-sorting framework, the receptor and products are considered as the primary solute. Thus the partition coefficient is defined as:

$$\varepsilon = \frac{[R_O]}{[R_A]} = \frac{[P_O]}{[P_A]}. \tag{14}$$

Furthermore, under the assumption of quasti-thermostatic partitioning, the volume fraction solute-rich phase must be variable given changing total synthesis and turnover of the respective solutes. We assume the concentration of solute in the aqueous and solute-rich phases are fixed, then the volume fraction of the solute-rich phase is proportional to the mean concentration of solutes within the cell. We thereby defined the “loss” of organelle $\lambda$ as a constant:

$$\lambda = 1 - \frac{[x_O]\phi_R}{\langle [x] \rangle} = \frac{\phi_A}{\phi_A + \varepsilon\phi_R} = \text{constant, if } \phi_R > 0. \tag{15}$$

1.2.3. Dynamics of the system and steady-state solutions

The dynamics of ligand-sorting system can be described by the following set of ODE:

$$\frac{d}{dt}\langle[x]\rangle=\beta_R\phi_A-\alpha_R[x_A]\phi_A-\delta[x_O]\phi_R,\ \text{x = R, RL, or P}$$

$$\frac{d}{dt}[L_A]=\beta_L-\alpha_R[L_A]-k[R_A][L_A]+k'[RL_A]-\frac{J_L}{V_c\phi_A}$$

$$\frac{d}{dt}[L_O]=-k[R_O][L_O]+k'[RL_O]-\delta[L_O]$$

$$\frac{d}{dt}\langle[RL]\rangle=\phi_A\left(k[R_A][L_A]-k'[RL_A]-\alpha_R[RL_A]-k_c[RL_A]\right)+J_L/V_c$$
$$+\phi_R\left(k[R_O][L_O]-k'[RL_O]-\delta[RL_O]-k_c[RL_A]\right)$$

$$\frac{d}{dt}\langle[P]\rangle=k_c\langle[RL]\rangle-\alpha_P[P_A]\phi_A-\delta[P_O]\phi_R \tag{16}$$

where $\langle[X]\rangle = \phi_A[X_A] + \phi_R[X_O]$ represents the weighted-average concentration of species X, and $V_c$ is the volume of cells to normalize the intake currents.

At the steady state, the ligand concentration in the organelle is given by:

$$\frac{d}{dt}[L_O]=0\Rightarrow[L_O]^*=\frac{k'\varepsilon}{k\varepsilon[R_A]^*+\delta}[RL_A]^* . \tag{17}$$

The ligand concentration in the aqueous phase, and average concentration of products are given by:

$$\frac{d}{dt}[L_A]=\beta_L-\alpha_R[L_A]-\left(k[R_A][L_A]-k'[RL_A]\right)A^{-1}=0$$

$$\frac{d}{dt}\langle[RL]\rangle=\phi_A\left(k[R_A][L_A]-k'[RL_A]\right)A^{-1}-\phi_A\tilde{\alpha}_{RL}[RL_A]=0 \tag{18}$$

where, to simply the equation, we defined extra a diffusion-limited scale-factor $A$ regarding intake current and apparent turnover of ligand-receptor complexes:

$$\tilde{\alpha}_{RL}=\alpha_{RL}+k_c+\frac{\phi_R}{\phi_A}\delta\left(\varepsilon+\frac{k'\varepsilon}{\delta+k\varepsilon[R_A]}\right), \tag{19}$$

$$A=\left(1+\frac{1}{V_c\phi_A}\frac{a}{4r_R}\frac{\varepsilon}{[R_O]+K_L}\right)^{-1} . \tag{20}$$

Then, the steady-state solutions can be given by Michaelis–Menten like equations:

$$[RL_A]^*=\frac{\beta_L}{\tilde{\alpha}_R}\frac{[R_A]^*}{[R_A]^*+K_m}$$

$$[L_A]^*=\frac{\beta_L}{\alpha_L}\frac{K_m}{[R_A]^*+K_m}$$

$$[P_A]=\frac{k_c}{\tilde{\alpha}_P}\left(\phi_A+\varepsilon\phi_R\right)\frac{\beta_L}{\tilde{\alpha}_R}\frac{[R_A]^*}{[R_A]^*+K_m} \tag{21}$$

where a Michaelis constant $K_m$, and apparent turnover of products is given by:

$$K_m = \frac{\alpha_L}{k} A + K_d \frac{\alpha_L}{\tilde{\alpha}_{RL}}$$

$$\tilde{\alpha}_P = \alpha_P + \frac{\phi_R}{\phi_A} \delta\varepsilon \quad . \tag{22}$$

in which $K_d = k'/k$ is the dissociation constant of the ligand-receptor complex.

1.2.4. Approximation of ligand sorting process under simplified conditions.

We assume all solutes are stable against degradation. Turnover of solutes are mainly governed by dilution rate caused by exponential growth. Also, if organelles have a constant volume fraction on average, then: $\alpha_R = \alpha_L = \alpha_{RL} = \alpha_P = \delta := \alpha$. Also, the synthesis rates of receptors and ligands are defined as the same: $\beta_R = \beta_L = \beta/2$.

If the irreversible step $k_c$ is significantly slower than the reversible steps $k$ and $k'$, we can apply steady-state approximation, and measure the apparent binding rate of ligand and receptors ($v_O$) in the system, molecules per unit time per unit volume by:

$$v_O = \alpha( \langle [RL] \rangle + \langle [P] \rangle) \quad . \tag{23}$$

If we assume $\alpha \gg k'\varepsilon$, which means $[L_O]$ is negligible in the system, then:

$$\tilde{\alpha}_{RL} = \alpha + k_c + \frac{\phi_R}{\phi_A} \alpha \left( \varepsilon + \frac{k'\varepsilon}{\delta + k\varepsilon [R_A]} \right) \cong \frac{\alpha}{\lambda} + k_c$$

$$K_m = \frac{\alpha}{k} A + K_d \lambda \tag{24}$$

Assuming that organelles have a strong contribution to the system, that the receptors are sparse, and that the exact volume of solute-rich phase is given by: $V_c \phi_R = VN$, we can further reduce $A$:

$$A = \left( 1 + \frac{1}{V_c \phi_A} \frac{a}{4r_R} \frac{\varepsilon}{[R_O] + K_L} \right)^{-1}$$

$$\cong \frac{4V_c \phi_A r_R \left( [R_O] + K_L \right)}{a\varepsilon}$$

$$\cong \frac{4V_c \phi_A r_R K_L}{a\varepsilon} = \frac{4\pi r^2 V_c \phi_A}{\varepsilon V a^2} = \frac{4\pi \phi_A}{\varepsilon \phi_R} \quad . \tag{25}$$

At the steady-state approximation, concentrations of ligands, and ligand-receptor complexes can be rewritten as:

$$\left[RL_A\right]=\frac{\beta}{2\phi_A}\frac{\lambda}{\alpha+\lambda k_c}\frac{\left[R_A\right]}{\left[R_A\right]+K_m};$$

$$\left[RL_O\right]=\frac{\varepsilon\beta}{2\phi_A}\frac{\lambda}{\alpha+\lambda k_c}\frac{\left[R_A\right]}{\left[R_A\right]+K_m}$$

$$\left[L_A\right]=\frac{\beta}{2\phi_A\alpha}\frac{K_m}{\left[R_A\right]+K_m};$$

$$\left[L_O\right]\cong 0 \quad . \tag{26}$$

And the average concentrations are given by:

$$\langle[RL]\rangle=\left[RL_A\right]\phi_A+\left[RL_O\right]\phi_R=\left[RL_A\right]\left(\phi_A+\varepsilon\phi_R\right)$$

$$=\frac{\beta}{2}\frac{1}{\alpha+\lambda k_c}\frac{\left[R_A\right]}{\left[R_A\right]+K_m}$$

$$\langle[L]\rangle=\left[L_A\right]\phi_A+\left[L_O\right]\phi_R=\frac{\beta}{2\alpha}\frac{K_m}{\left[R_A\right]+K_m} \tag{27}$$

The apparent binding rate of ligand sorting by organelles ($v_O$) is given by:

$$v_O=\frac{\beta}{2}\frac{\left[R_A\right]}{\left[R_A\right]+K_m} \, . \tag{28}$$

The partition of ligands is defined as:

$$\varepsilon_L=\frac{\left[L_O\right]+\left[RL_O\right]+\left[P_O\right]}{\left[L_A\right]+\left[RL_A\right]+\left[P_A\right]}$$

$$=\frac{\varepsilon\lambda\left[R_A\right]}{K_m+\lambda\left[R_A\right]}$$

$$=\frac{\varepsilon\left[R_A\right]}{\frac{4\pi\phi_A}{\lambda\varepsilon\phi_R}\frac{\alpha}{k}+K_d+\left[R_A\right]}$$

$$=\frac{\varepsilon\left[R_A\right]}{\frac{4\pi}{1-\lambda}\frac{\alpha}{k}+K_d+\left[R_A\right]} \quad . \tag{29}$$

If the free, unrecruited receptors always have a small concentration, by defining $\eta = 1\text{-}\lambda$, an upper-limit approximation of the partition of ligand can be given by:

$$\varepsilon_L\leqslant\frac{\varepsilon\left[R_A\right]}{\frac{4\pi}{\eta}\frac{\alpha}{k}+K_d}=\frac{\left[R_O\right]}{\frac{4\pi}{\eta}\frac{\alpha}{k}+K_d} \, . \tag{30}$$

Considering the concentration of aqueous receptors is a proportion to the average concentration of receptors within the cells:

$$\lambda\langle[R]\rangle = \phi_A[R_O], \tag{29}$$

the apparent binding rate can be rewritten as:

$$v_O = \frac{\beta}{2}\frac{\langle[R]\rangle}{\langle[R]\rangle + K_M}, \tag{30}$$

where the adjusted M-M constant $K_M$ is given by:

$$K_M = \left(\frac{\alpha}{k}A + K_d\lambda\right)\frac{\phi_A}{\lambda} = \phi_A\left(\frac{4\pi}{1-\lambda}\frac{\alpha}{k} + K_d\right). \tag{31}$$

If at the steady state, the sum of the concentrations of receptors, ligand-receptor complexes, and products is equal to $\beta/\alpha$, we can solve the concentration of receptors:

$$\langle[R]\rangle + \langle[RL]\rangle + \langle[P]\rangle = \frac{\beta}{2\alpha}$$

$$\Rightarrow \langle[R]\rangle = \frac{1}{2}\left(\sqrt{K_M\left(K_M + 2\frac{\beta}{\alpha}\right)} - K_M\right)$$

$$\Rightarrow [R_A] = \frac{\lambda}{\phi_A}\frac{1}{2}\left(\sqrt{K_M\left(K_M + 2\frac{\beta}{\alpha}\right)} - K_M\right). \tag{32}$$

Therefore, instead of being maintained in a fixed concentration (*11*), the steady-state concentration of free receptors in aqueous phase may be changed due to ligand-receptor binding, consistent with recent discovery (*33*).

### 1.3. Comparison between well-mixed system and ligand-sorting system across changing synthesis rates

We first summarize the steady state solutions of apparent binding rates. The solution of the well-mixed reactions can be given by Eq. 7, and that of the ligand-sorting system can be further reduced from Eq. 30 and 32. We reformed them into similar formula as the following:

$$v_M = \left(\frac{\beta}{\sqrt{\alpha^2/k_i + 2\beta} + \sqrt{\alpha^2/k_i}}\right)^2$$

$$v_O = \left(\frac{\beta}{\sqrt{\alpha K_M + 2\beta} + \sqrt{\alpha K_M}}\right)^2 \tag{33}$$

To unify our results in two systems, we define a "generic" apparent binding rates ($v$) as a function of total synthesis rate ($\beta$), and "resistance" of the system to associate ligands and receptors ($\rho$):

$$v(\rho, \beta) = \left(\frac{\beta}{\sqrt{\rho + 2\beta} + \sqrt{\rho}}\right)^2. \tag{34}$$

Apparently higher the $\rho$ is, less flux of synthesis events will turn into ligand-receptor binding events, where resistance for the well-mixed and ligand-sorting systems are given by:

$$\rho_M = \frac{\alpha^2}{k_i} \cong \frac{\alpha^2 k'}{k(k_c + \alpha)}$$

$$\rho_O = \alpha K_M \cong \alpha\phi_A\left(\frac{4\pi}{\eta}\frac{\alpha}{k} + K_d\right). \tag{35}$$

Next, we tested the parameter sensitivity of the two systems by measuring the partial derivative of generic binding rate. We particularly focused on the total synthesis rate ($\beta$) that can be easily altered by changing the induction levels in experiments. The sensitivity to total synthesis rate is given by:

$$\frac{\partial v(\rho,\beta)}{\partial \beta} = \frac{1}{2}\left(1 - \frac{\sqrt{\rho}}{\sqrt{\rho + 2\beta}}\right) = \sqrt{\frac{v}{\rho + 2\beta}}$$

$$\Rightarrow \lim_{\rho \to \infty} \frac{\partial v(\rho,\beta)}{\partial \beta} = 0 \quad . \qquad (36)$$

Thus, higher the $\rho$ and the lower $\beta$ is, the system is less sensitive to the change of synthesis rate. This suggests that lower the $v$ is, more robust the system is. For $\rho$ converging to the infinity, the system is in absolute robustness.

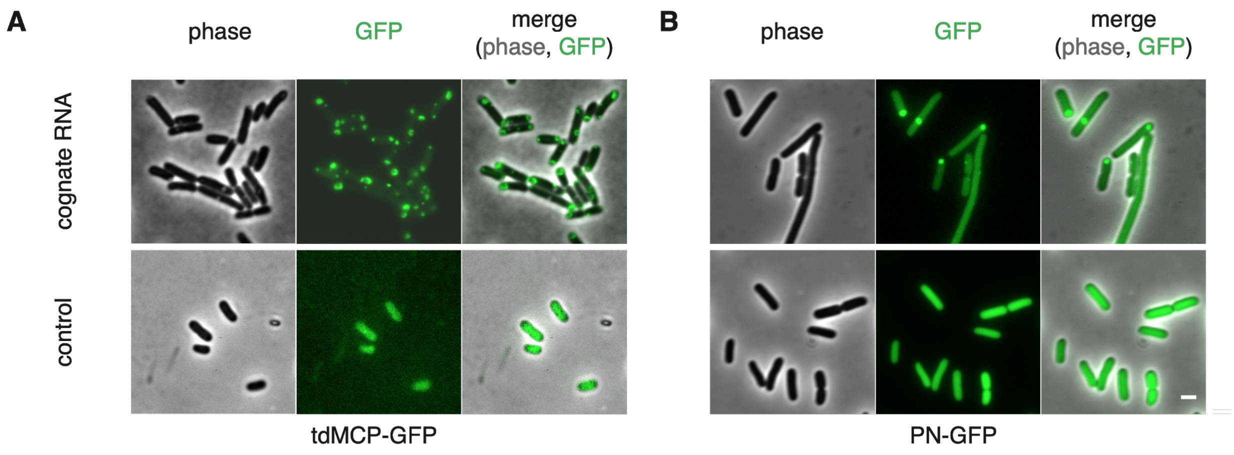


**Fig. S1. Spatial distribution of ABD-GFP with or without cognate rCAG-aptamers.**

Phase contrast and fluorescent imaging of cells expressing cognate pairs of rCAG-aptamer and ABD-GFP, or ABD-GFP alone. (A) tdMCP-GFP, and (B) PN-GFP. Shown from top to down are cells expressing rCAG-aptamer that displayed fluorescence foci, and cells expressing ABD-GFP alone that displayed homogeneous distribution of fluorescence. Scale bar, 1 μm.

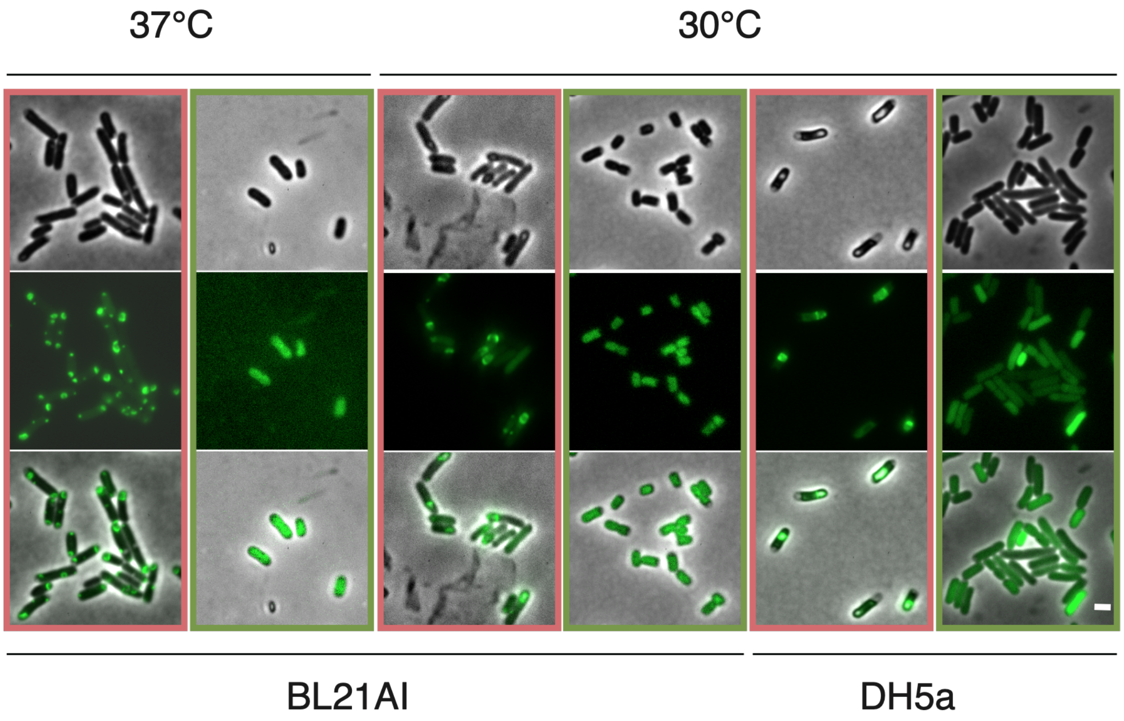


**Fig. S2. RNA phase condensates validated under different contexts.**

Phase contrast and fluorescent imaging of cells expressing tdMCP-GFP with (red boxes) or without (green boxes) rCAG-MS2 in BL21AI or DH5a, at 30 or 37°C. Under all contexts, cells expressing rCAG-MS2 formed fluorescent foci, and cells expressing only tdMCP-GFP displayed homogeneous fluorescence. Scale bar, 1 µm.

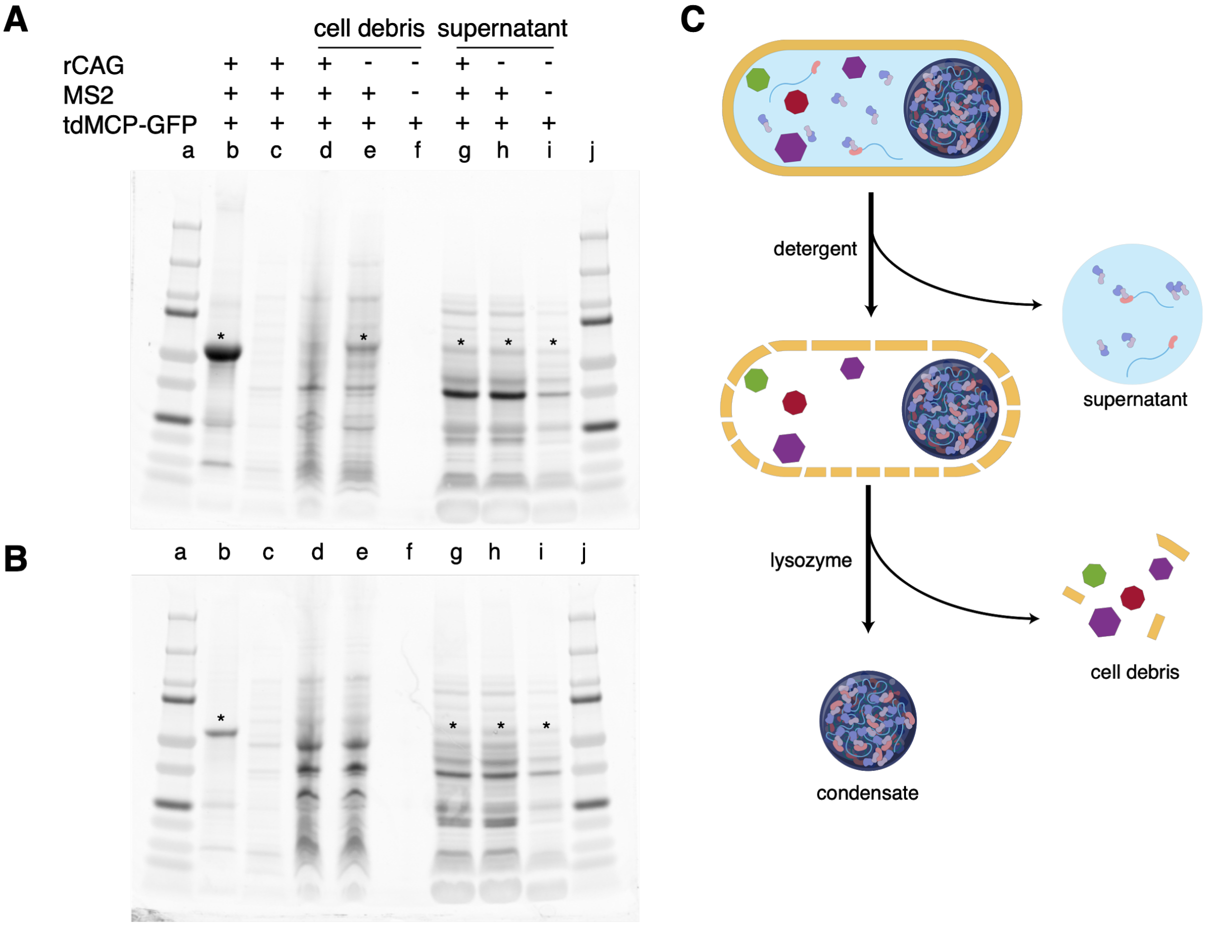


**Fig. S3. Purification of RNA condensates.**

(A and B) SDS-PAGE of condensate purification assay. Indicated systems were expressed in BL21AI (A) or DH5a (B). (a and j) protein ladders, (b) condensate purified from cells expressing rCAG-MS2 and tdMCP, (c) washed remained cell debris, (d - f) cell debris from indicated systems, (g - i) supernatant from indicated systems. * highlighted the bands of complete tdMCP-GFP, judged by molecular mass. (C) Schematic of condensate purification assay (also see Material and Methods). Harvested cells were sequentially treated by detergent, and lysozyme, to isolate supernatant, large components of cell debris, and condensate of interest.

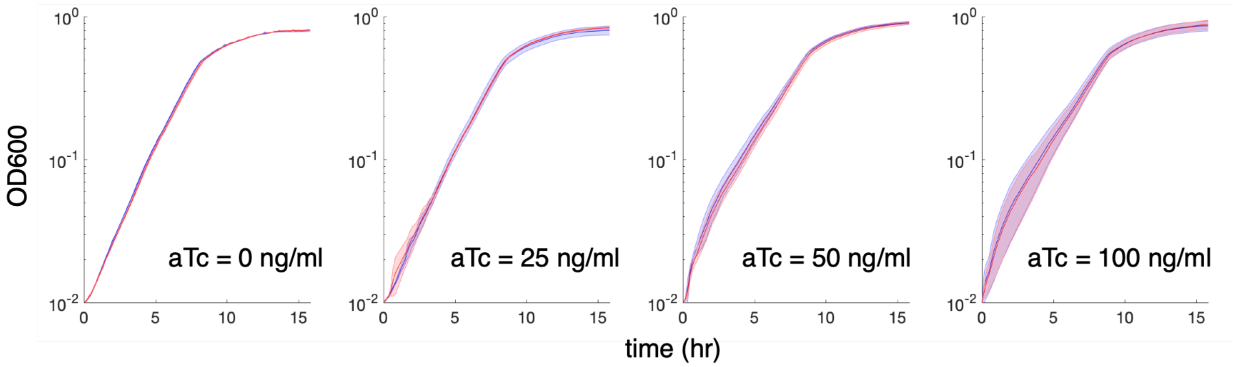


**Fig. S4. Comparison of bacterial growth with or without rCAG.**

Growth curves of cells expressing either tandem 12 MS2 aptamer without CAG repeats (red), or rCAG-MS2 (blue) measured in 96-well plate by plate readers overnight. From left to right are 4 different aTc induction levels, 0, 25, 50, and 100 ng/ml. Coefficients of determination (Rsq) were calculated to fit growth curves with rCAG-MS2 versus with simple MS2 aptamers rCAG-MS2 by identity line as 0.9998, 0.9974, 0.995, and 0.9987 for aTc inductions from 0 to 100 ng/ml, suggesting that cells expressing MS2 aptamers with or without CAG repeats had exactly the same growth under the same aTc induction level. Shaded error bars, standard deviation from 3 technical replicates.

A B

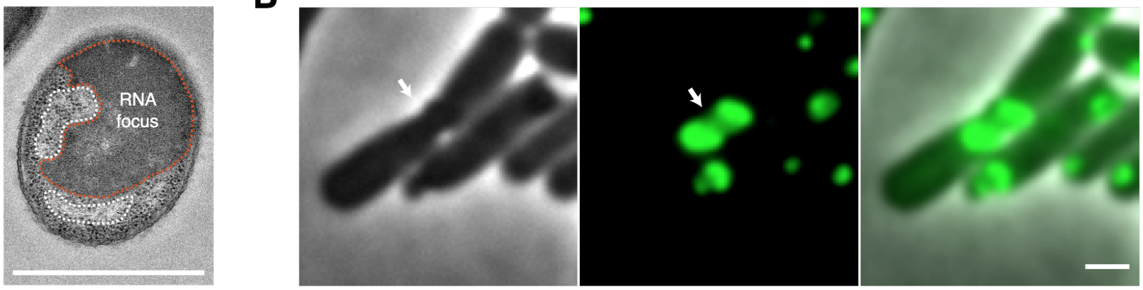


**Fig. S5. Irregular shapes of RNA foci contacted with subcellular structures.**

(A) TEM image of RNA focus contacting nucleoid. The nucleoid area (white dash line) is brighter, and RNA focus (red dash line) is denser, compared to the unlabeled cytosolic region. (B) Shown from left to right are phase, GFP (highlighted fluorescent foci), and overlay images of RNA focus that were reshaped by a division ring. The position of division is highlighted by the arrow. Scale bar, 1 μm.

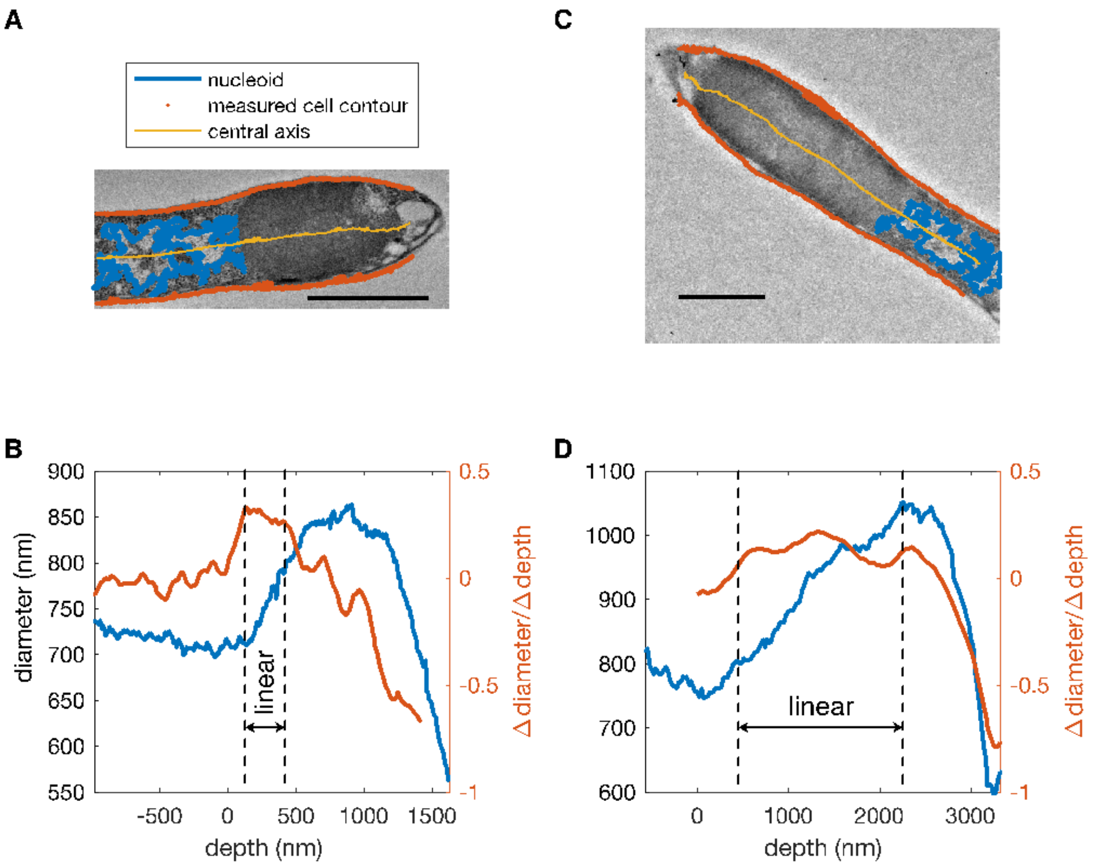


**Fig. S6. Surface tension of TEARS enlarges the diameter of cells.**

(A and C) Processed TEM image of two representative large TEARS adhered to the cellular membrane. Nucleoids, cellular contour and cellular axes are labeled in blue, red and yellow, respectively. The cellular axes are plotted by the midpoints of cellular contours. Scale bar, 1 µm. (B and D) Cell diameter change in response to depth of TEAR droplets along the cell axis (blue), and derivative of cell diameter with droplets depth (red) measured as the slope of a linear regression with a sliding window of 25 data points (~ 206 nm in B, and ~ 577 nm in D). The linear regimes of cellular diameter increase with depth are highlighted. For each pair of measured positions on the cellular contours, the respective depth of droplets is defined as the distance between midpoints of measured positions and the point of intersection between cellular axes and droplet surface.

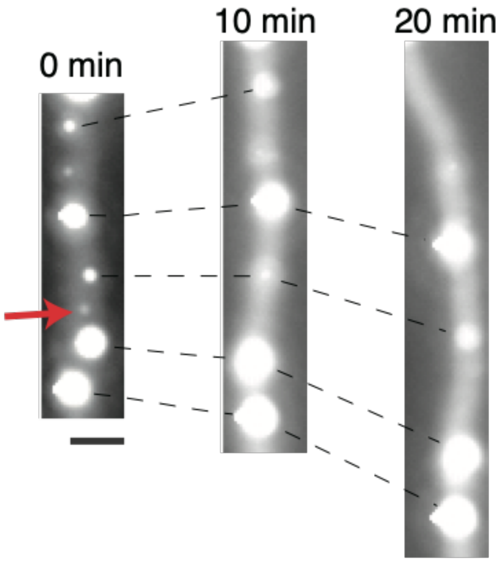


**Fig. S7. Ostwald ripening.**

Presence of large and small TEARS droplets within the same cell leads to growth of the former (black lines), and dissipation of the latter (red arrow). Scale bar, 1 μm.

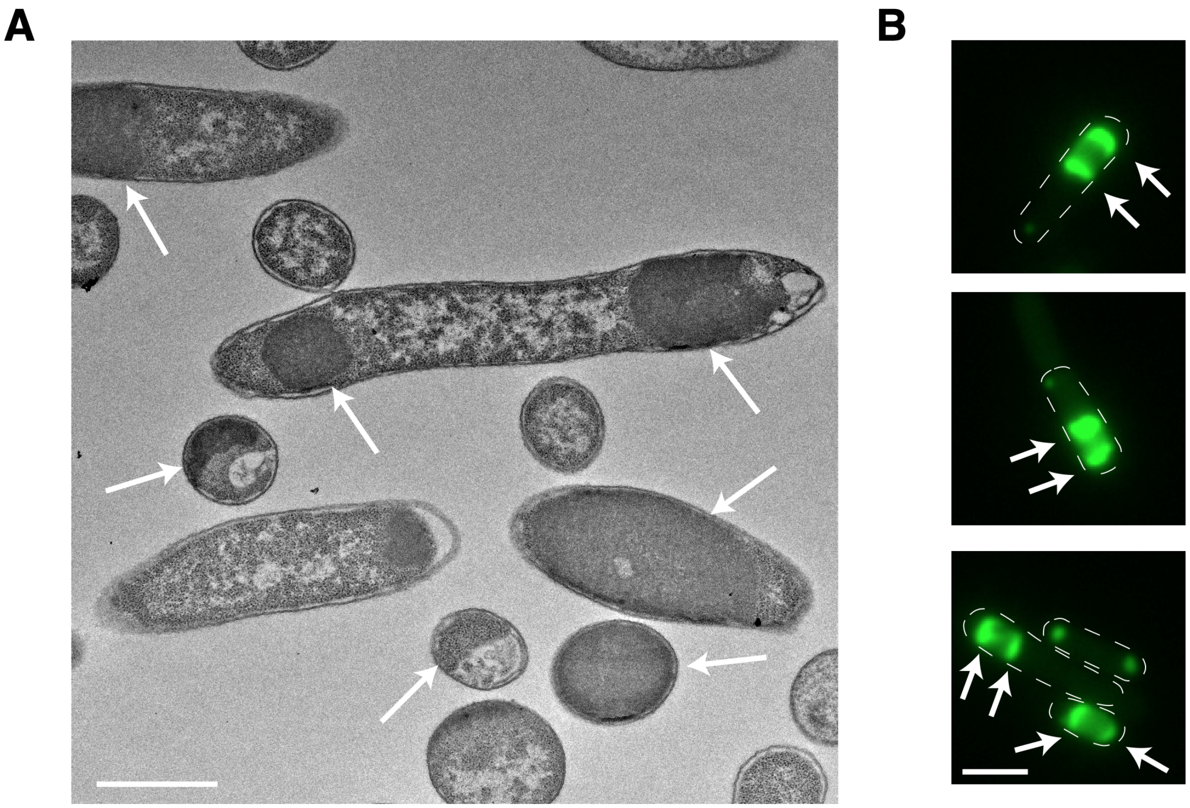


**Fig. S8. Multiple layers of TEARS composed of rCAG-MS2 and tdMCP-GFP**

(A) TEM image of cells expressing rCAG-MS2 with tdMCP-GFP. Under coexpression of rCAG-MS2 and tdMCP-GFP, all TEARS are material dense, showing that no droplets are void of RNAs in the middle of them. (B) Additional representative examples of TEAR droplets with outer GFP-rich layer and inner GFP-poor core. Arrows highlight the GFP-rich layers. Dashed lines show manual segmentation of cell contours. Scale bar 1 μm.

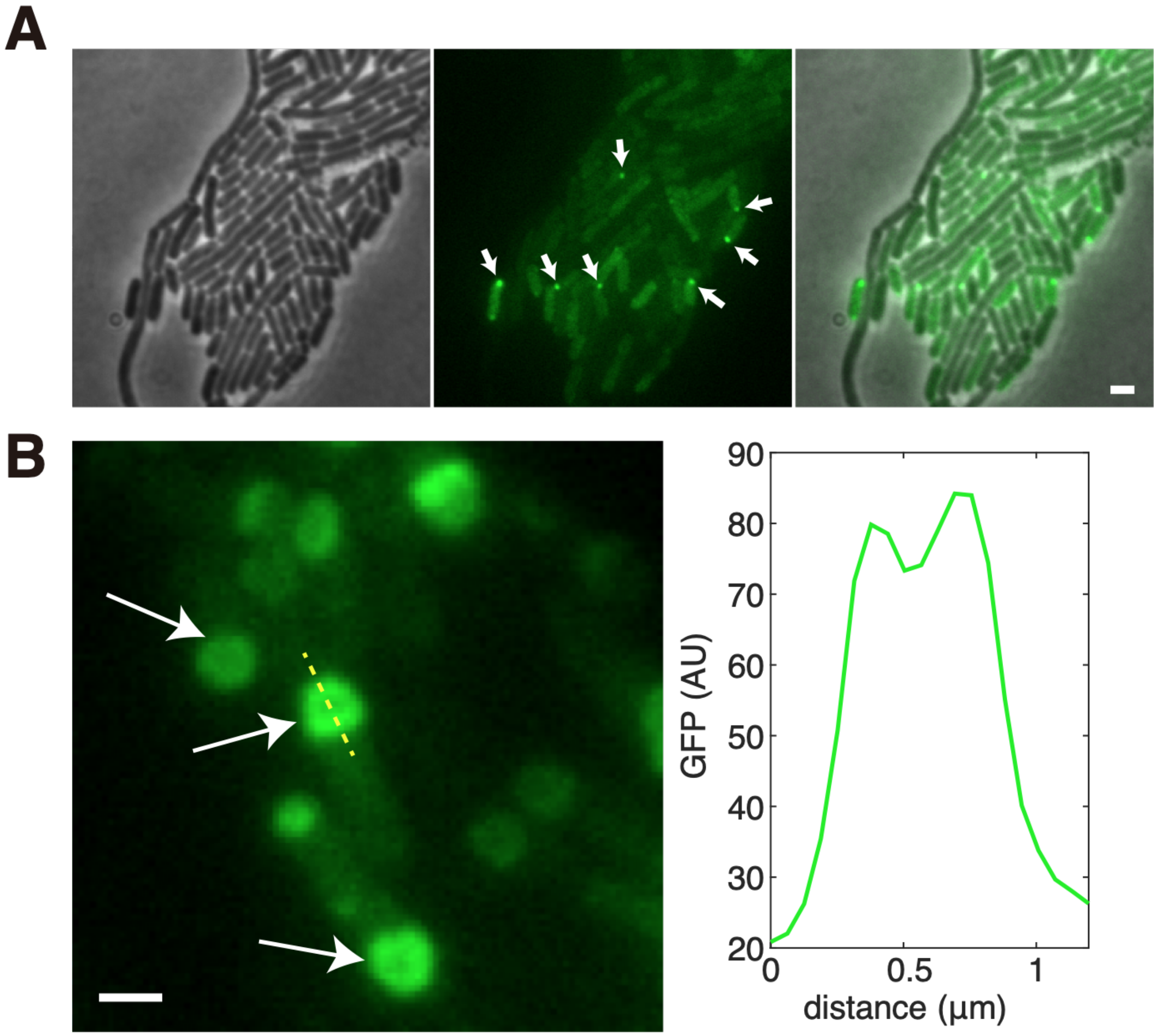


**Fig. S9. FA-FB associated in the TEAR droplets versus spontaneous FA-FB aggregates.**

(A) In cells devoid of RNA droplets, expressing FA-tdMCP and FB,random oligomerization occasionally forms aggregates with high fluorescence, highlighted by arrows. Shown from left to right are phase, GFP, and overlay. (B) In comparison, in presence of RNA droplets, FA and FB organized droplets showed a signature ring-like multilayered structure. Shown from left to right are magnified images of representative cells in the GFP channel (arrows highlighted droplets with dimmer centers), and line profile (yellow line for GFP profile). The structure is similar to observations shown in Fig. 3C, indicating the liquid phase. No such forms were observed in absence of RNA droplets. Scale bar, 1 µm.

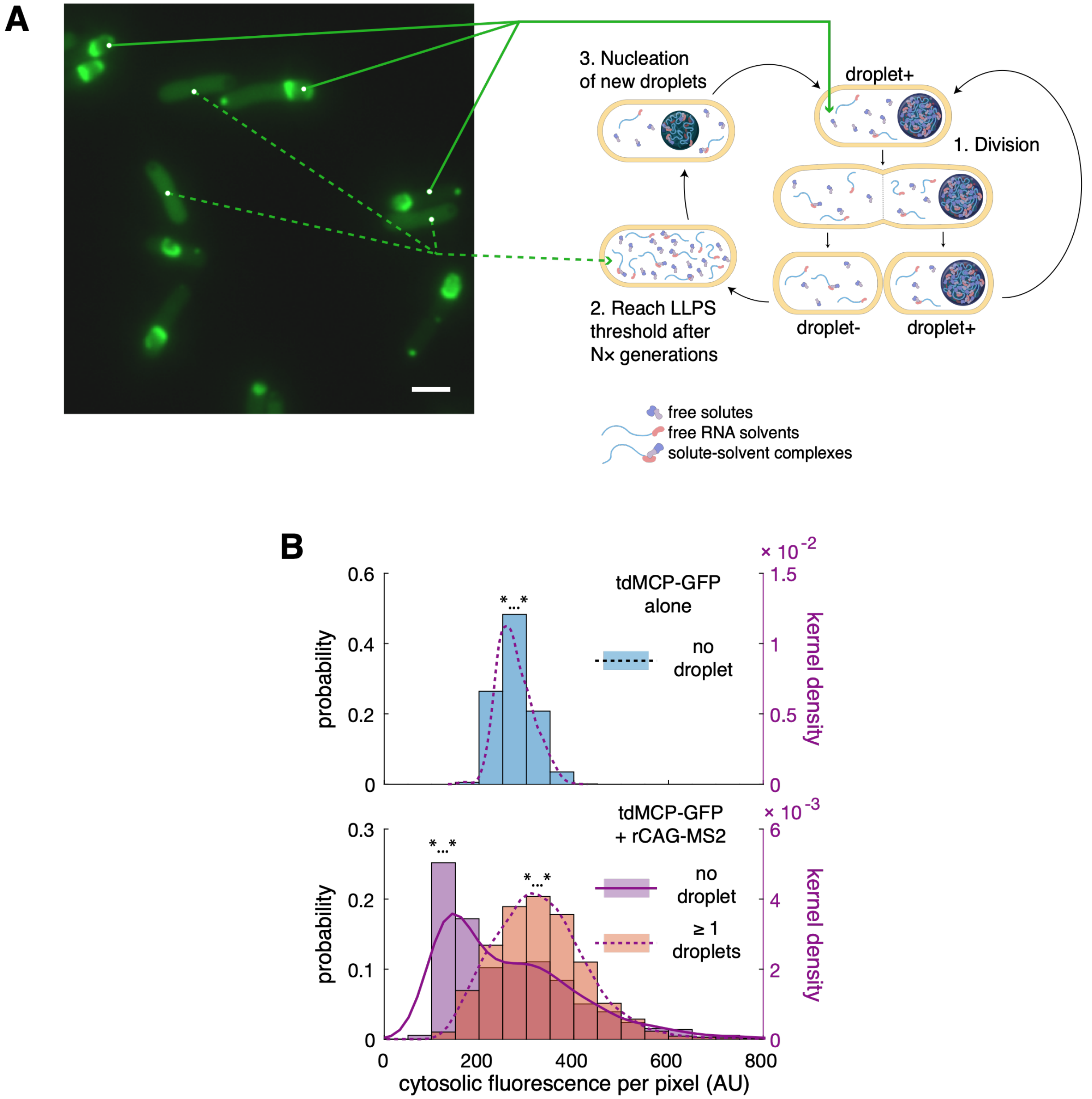


**Fig. S10. Variability induced by asymmetric division of TEARS.**

(A) Division of cells carrying droplets generates two types of daughter cells, cells that inherit the droplets from mother cells (droplet +), and cells that lose the droplet (droplet -). Cells without droplets can accumulate RNA solvents and solutes for several generations, and generate new droplets from scratch. Shown from left to right, GFP images, and diagram of the asymmetric division. Green arrows link three representative cells each from two subpopulations to the diagram. Scale bar is 1 µm. (B) On the top, distributions of cytosolic mean fluorescence intensity (left y-axis histogram bars, and right y-axis kernel density) for control cells expressing tdMCP-GFP alone (top, blue bars, and magenta dashed lines). At the bottom, cells expressing tdMCP-GFP and rCAG-MS2 (bottom) differentiates into two subpopulations due to asymmetric division, one without droplets (purple bars, solid line), and one with over 1 droplets (orange bars, dashed lines).

Compared to controls, both subpopulations have a wider range of fluorescence, and in addition, the subpopulation lost droplets have lower fluorescence, and the one containing droplets have higher fluorescence, on average, which further contributes to the variability of the entire population. *...*, all three distributions are distinct from each other, p-value $< 10^{-70}$, by pairwise Kolmogorov–Smirnov test. AU, arbitrary units.

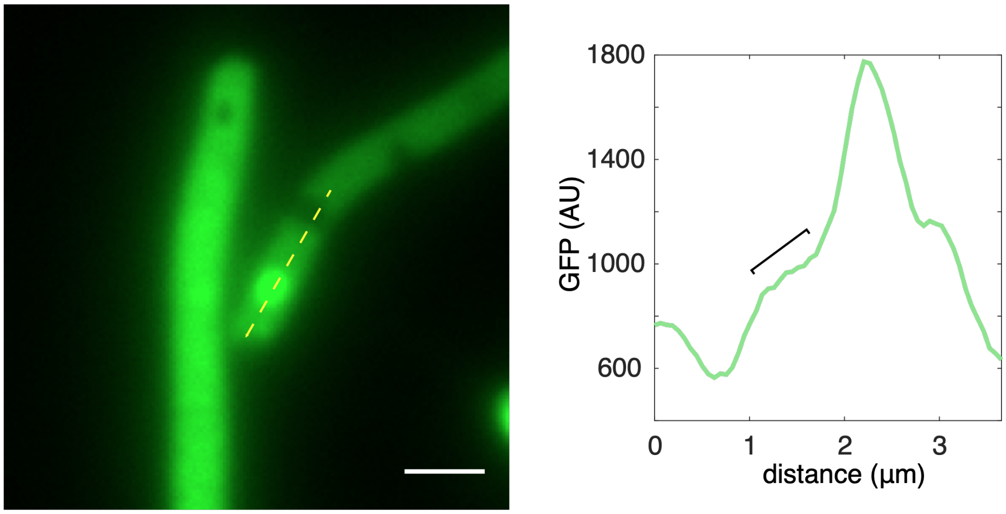


**Fig. S11. Other examples of intracellular asymmetry of solutes.**

Similar to Figure 3C, another representative example of coexistence of foci enriched GFP and void of GFP with no contact, in cells expressing BoxB-based systems. Shown from left to right are GFP image and line profiles for GFP channel (yellow dashed line, GFP profiles). A gradient of GFP exists between GFP-rich and GFP-poor phases, highlighted by black line. AU, arbitrary units. Scale bar, 1 μm.

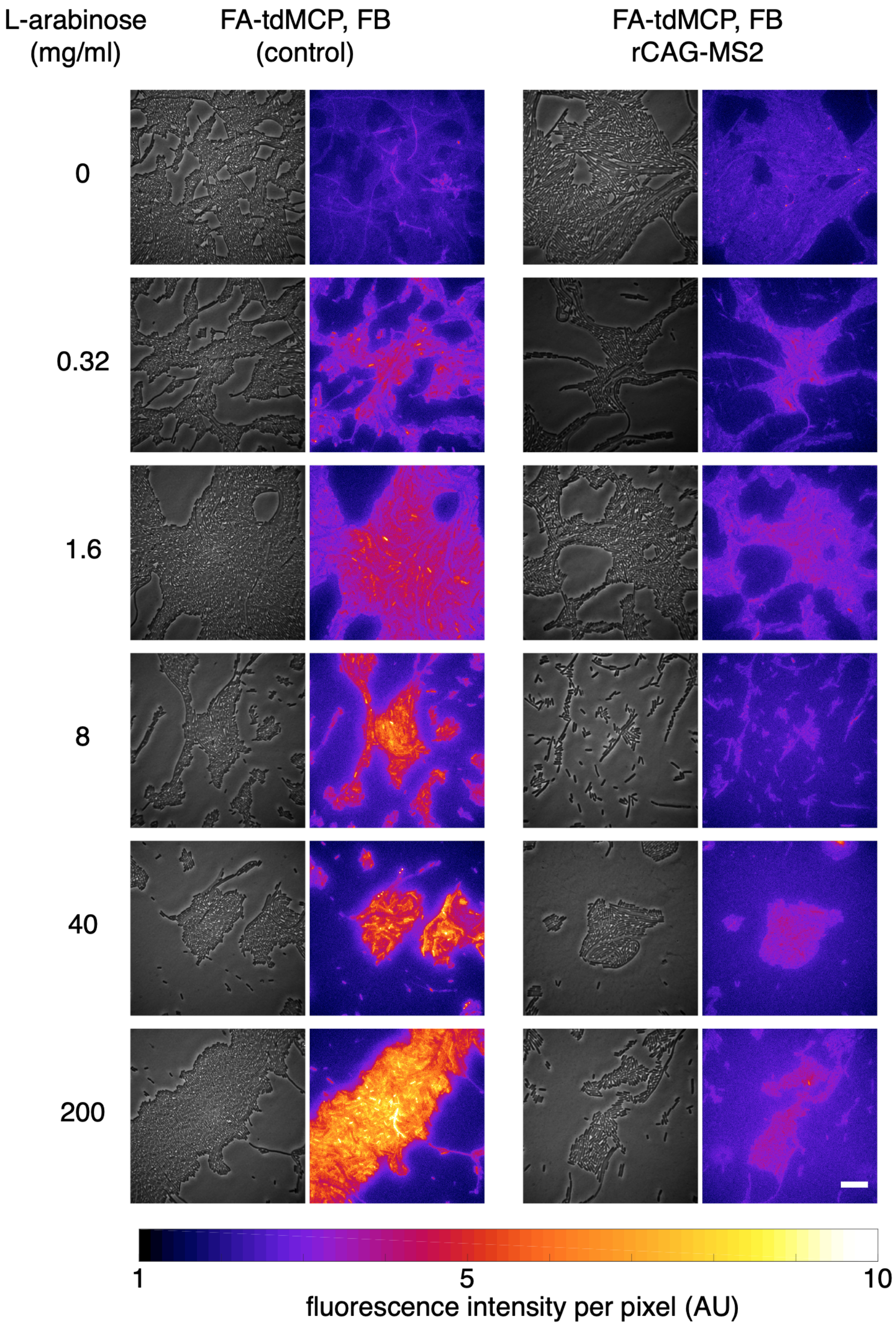


**Fig. S12. BiFC with or without ligand sorting across changing synthesis rates.**

Images of representative exponential-phase cells expressing FA-tdMCP, FB under indicated L-arabinose inductions, with (left) or without (right) fully-induced rCAG-MS2. Shown from top to bottom, the arabinose increased from 0% to 0.2%. For each strain under indicated induction, shown from left to right are phase contrast image, and GFP fluorescence intensity map. Scale bar, 10 μm. Colormap span from $10^3$ to $10^4$ arbitrary units (AU).

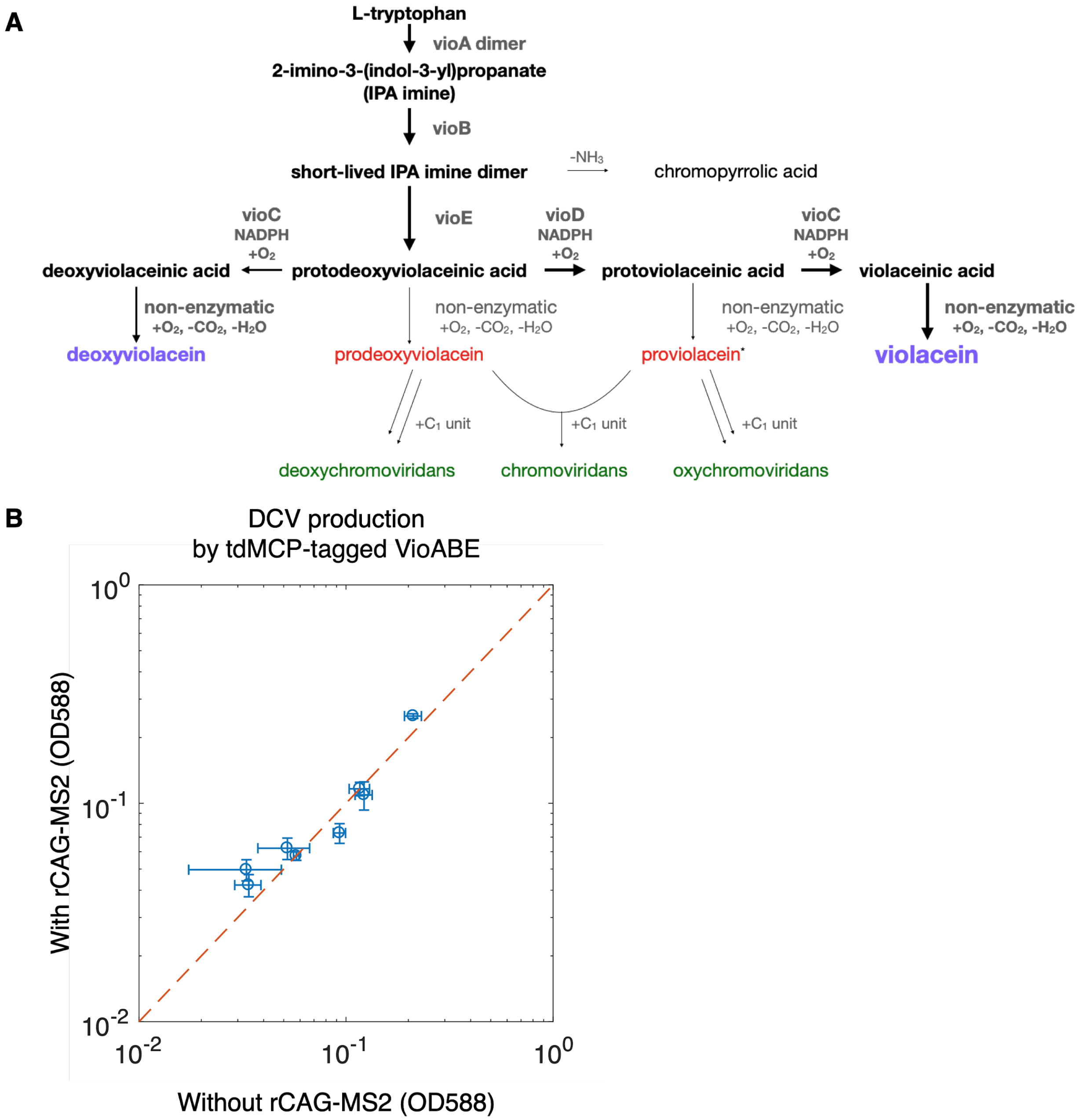


**Fig. S13. Supplementary material for deoxyviolacein assay**

(A) Diagram of complete pathway of Vio operon (26). Amino acid L-tryptophan is first catalyzed by VioA dimer into IPA imine. VioB dimierizes IPA imine into an unstable intermediate that can turn into color-less chromopyrrolic acid by spontaneous deamination. VioE catalyzes IPA imine dimer into protodeoxyviolacein acid (PDA), where the branches starts. With only VioC, PDA can be processed by VioC into deoxyviolaceinic acid. In the presence of both VioC and VioD, the majority of PDA will be oxidized into protoviolaceinic acid, which can be further processed by VioC into violaceinic acid. Violaceinic acid and other three types of protodeoxy-, proto-, deoxy-violaceinic acid can be spontaneously oxidized, and decarboxylated into respective analogs of violacein. Violacein and deoxyviolacein are violet-color pigments, yet prodeoxyviolacin and proviolacein are red-color pigments, and can further homo-, or hetero-dimerize into green-color deoxychromoviridans, chromoviridans and oxychromoviridans with one more carbon added (+C1 unit), through an unclear mechanism. In the natural strain containing VioABCDE, the majority of the flux goes into violacein production, and a minority goes into deoxyviolacein. However, in the absence of VioD, the metabolic direction towards deoxychromoviridans can be favored. (B). Clustering of VioABE has an insignificant influence on production of DCV. Coexpressing tdMCP-tagged VioABE with or without rCAG-MS2 produced similar amounts of DCV, regardless of promoter strengths. Production of DCV is measured by OD at 588 nm using a plate reader. Promoters were randomly mutated from BBa_J23105 that was used to drive express VioABE in other experiments. Error bars are standard deviation from 5 biological replicates. Dash red lines show the diagonal.

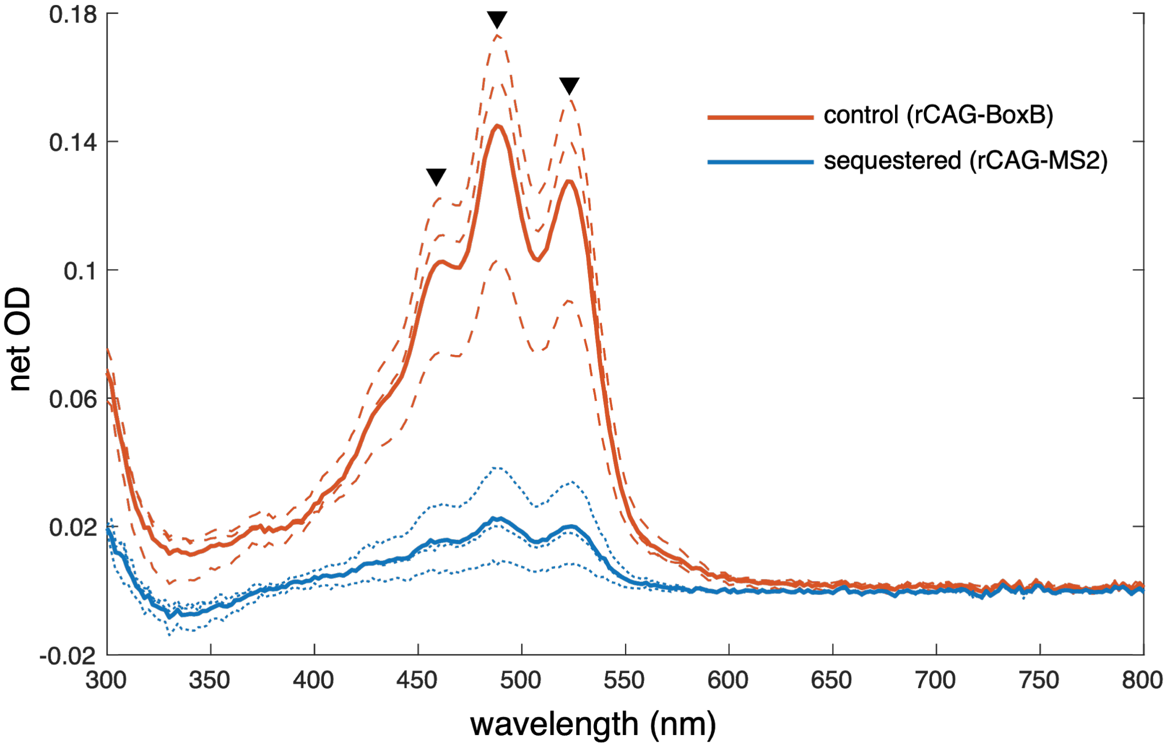


**Fig. S14. Absorption spectrum of extracts of lycopene production.**

Products of cells expressing untagged CrtE, tdMCP-tagged CrtBI, and cognate TEAR droplets (rCAG-MS2, blue), or orthogonal droplets (rCAG-BoxB, red) were extracted by DMSO. Absorption of products were scanned from 300 nm to 800 nm. Background signal of extracts of DH5a cells expressing no lycopene pathway was subtracted. Dash lines show individual absorption spectrums of 3 biological replicates each, and solid lines show the mean absorption. Black triangles highlight absorption maxima of lycopene-DMSO solution, 462, 488, and 522 nm.

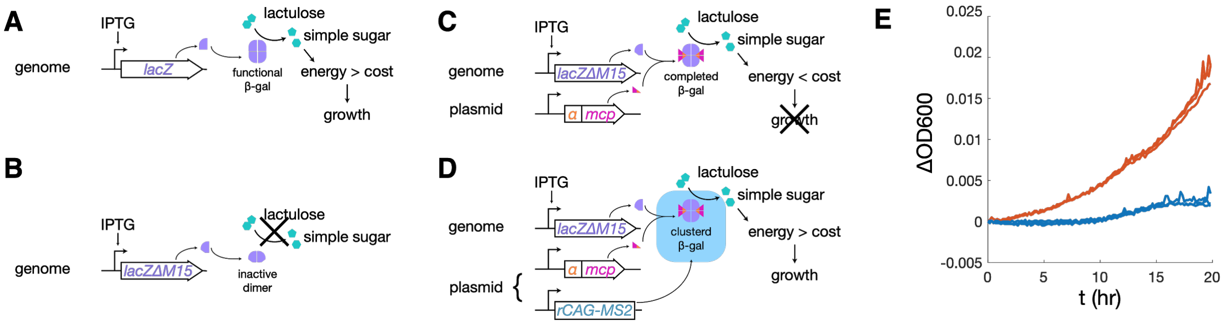


**Fig. S15. Supplementary material for α-complementation assay.**

(A-D) Schematic diagrams for detailed mechanisms. (A) Wild-type strains are capable of processing lactulose under enough IPTG induction to support growth by functional tetrameric β-gal encoded in genomic *lacZ*. (B) The genome of M15 mutants only encodes a fragment (ω-peptide) of β-gal that forms into a non-functional dimer, thereby unable to process lactulose. (C) The α-peptides encoded by *lacZα* can associate with ω-peptides to complete into a functional β-gal tetramer. Thus M15 mutants expressing *lacZα*, are capable of processing lactulose. However, the complemented β-gal only has a low efficiency that produces energy lower than the cost of growth, which does not support growth. (D) M15 mutants expressing TEAR droplets to cluster α-complementation are capable of processing lactulose in a high efficiency that supports growth. (E) Growth curves of DH5a cells expressing ω-fragment and α-tdMCP alone for control (red), or with rCAG-MS2 for clustering (blue), with 3 replicates each. (F) Agar assay of DH5a cells expressing endogenous ω-fragment (25 μM IPTG induced), α-tdMCP, with rCAG-MS2 (+) or rCAG-BoxB (-), imaged by FITC channel of autofluorescence. (E) α-tdMCP alone cannot rescue the growth defect of DH5a on lactulose. Growth curves of DH5a cells expressing ω-fragment and α-tdMCP alone (red), or with rCAG-MS2 for clustering (blue), with 3 replicates each. The initial OD values were subtracted.

**Table S1. Genetic part synthesized in this study.**

(a) pHR-Tre3G-47xCAG-12xMS2 was a gift from Ron Vale (Addgene plasmid # 99148 ; http://n2t.net/addgene:99148 ; RRID:Addgene_99148) (*21*).
(b) BBa_K1100100 is a plasmid provided by Haotian Guo (first author of this study).
(c) BBa_K274004 was designed by Shuna Gould and constructed by the iGEM 2009 Cambridge team. It was acquired from iGEM part registry through 2012 Distribution.

| Name | Source |
| --- | --- |
| 47×CAG, 12×MS2, WHP Posttranscriptional Response Element | PCR from pHR-Tre3G-47xCAG-12xMS2 (a) |
| 12×BoxB | annealed oligo |
| superfolder GFP | PCR from BBa_K1100100 (b) |
| PA1lacO-1, λN | gblock |
| BBa_B1004, BBa_J23105, Golden Gate assembly cloning sites | gblock |
| mKate-PT linker-ELK16, pFAB815 | assembled from gblock |
| vioA | PCR from BBa_K274004 (c) |
| vioB | PCR from BBa_K274004 |
| vioE | PCR from BBa_K274004 |
| BBa_B0034, vioC | gene fragment |
| riboJ, BBa_B0034, vioD | gene fragment |
| tdMCP_1 (tdMCP same-sense sequence 1) | gblock |
| tdMCP_2 | gblock |
| tdMCP_3, phA-1 terminator | gblock |
| tdMCP_4, thrL terminator | gblock |
| RcCrtA | gene fragment |
| EuCrtE, | gene fragment |
| riboJ, EuCrtB | gene fragment |
| PlmJ, EuCrtI | gene fragment |
| lacZα | gblock |
| BBa_R0040 | annealed oligo |

**Table S2. Designated backbones used for cloning.**

| Backbone | Information | Source |
|---|---|---|
| pSB1A3 | https://parts.igem.org/Part:pSB1A3 | iGEM part registry, 2017 Distribution |
| pSB4K5 | https://parts.igem.org/Part:pSB4K5 | iGEM part registry, 2017 Distribution |
| pCOLADuet-1 | https://www.emdmillipore.com/PR/en/product/pCOLADuet-1-DNA-Novagen,EMD_BIO-71406 | Novagen |
| pACTet | double terminator, p15A, CmR, TetR, PLtetO-1 | this study |
| pSB1A3-J23105 | pSB1A3, BBa_B1004, BBa_J23105, Golden Gate assembly cloning sites | this study |

**Table S3. Plasmids characterized in this study.**

| Name | Backbone | Insert | Source | Sequence |
|---|---|---|---|---|
| prCAG-MS2 | pACTet | 47×CAG, 12×MS2, WHP Posttranscriptional Response Element | this study | https://benchling.com/s/seq-c98PnzCCphq5eKL7P2GI |
| prCAG-BoxB | pACTet | 47×CAG, 12×BoxB | this study | https://benchling.com/s/seq-vSRHVWCpW7KH8KRKtLwG |
| pIG-K133 | pPROTET.E | P6×HN, tdMCP, GFPmut3 | (*45*) | |
| pPN-GFP | pSB4K5 | pA1lacO-1, λN, superfolder GFP | this study | https://benchling.com/s/seq-y03ghv9TXzybTHsRmVSS |
| pmKate-ELK16 | pSB1A3 | BBa_J23105, BBa_B0034, mKate-PT linker-ELK16, pFAB815 | this study | https://benchling.com/s/seq-jyAO3Ui2o1qaOvnSH1BT |
| pFA-MCP | pCDFDuet-1 | FA-tdMCP | (*6*) | |
| pFB | pCOLADuet-1 | FB-tandem dimer PP7 coat protein | (*6*) | |
| pVioAMBMEM | pCOLADuet-1 | BBa_B1004, BBa_J23105, BBa_B0034, vioA-tdMCP_1, BBa_B0034, vioB-tdMCP_2, BBa_B0034, vioE-tdMCP_3, pheA-1 terminator | this study | https://benchling.com/s/seq-uTxeY5EFtoSDO5qTu8RC |
| pVioCM | pSB1A3-J23105 | BBa_B0034, vioC-tdMCP_4, thrL terminator | this study | https://benchling.com/s/seq-2I7pwgAjNR4FhvRBgX3F |
| pVioCD53 | pSB1A3-J23105 | BBa_B0034, vioC, riboJ, B0034, vioDΔ(264-372), thrL terminator | this study | https://benchling.com/s/seq-KyBTdM1oAwGl9FvXtCos |

| | | | | |
|---|---|---|---|---|
| pCrtEBMIM | pSB1A3-J23105 | BBa_B0034, EuCrtE, riboJ, BBa_B0034, EuCrtB-tdMCP_2, PlmJ, BBa_B0034, EuCrtI-tdMCP_3, thrL terminator | this study | https://benchling.com/s/seq-IER5dBPXkLMpwvYAwUkZ |
| pCrtAEBMIM | pSB1A3-J23105 | BBa_B0034, RcCrtA, BBa_B0015, BBa_B1004, BBa_J23105, BBa_B0034, EuCrtE, riboJ, BBa_B0034, EuCrtB-tdMCP_2, PlmJ, BBa_B0034, EuCrtI-tdMCP_3, thrL terminator | this study | https://benchling.com/s/seq-ZEywoudl4dvuR5apO6Cx |
| placZα-tdMCP | pSB1A3 | BBa_R0040, BBa_B0034, lacZα-tdMCP_1 | this study | https://benchling.com/s/seq-oKrtIqEiL0q5JBqYMlqO |
| prCAG-RBS_library-sfGFP (a) | pACTet | 47×CAG, RBS library, superfolder GFP | this study | https://benchling.com/s/seq-H5FJq0MioMg964S635as |
| prCAG-RBS"wt"-sfGFP | pACTet | 47×CAG, RBS"wt", superfolder GFP | this study | https://benchling.com/s/seq-D4sYjBE4OzI56r2XoBel |
| prCAG-RBS"m1"-sfGFP | pACTet | 47×CAG, RBS"m1", superfolder GFP | this study | https://benchling.com/s/seq-WCf3ceK01oP9HeYqJvDa |
| prCAG-RBS"m2"-sfGFP | pACTet | 47×CAG, RBS"m2", superfolder GFP | this study | https://benchling.com/s/seq-NElJc6JeMPXFCuHHSPRT |
| prCAG-RBS"m3"-sfGFP | pACTet | 47×CAG, RBS"m3", superfolder GFP | this study | https://benchling.com/s/seq-buTUGcvWsteEhrEtJZvp |
| prCAG-RBS"m4"-sfGFP | pACTet | 47×CAG, RBS"m4", superfolder GFP | this study | https://benchling.com/s/seq- |

| | | | | |
|---|---|---|---|---|
| | | | | F5JUUWUNudRpgvF0AA3O |
| pRBS"wt"-sfGFP | pACTet | RBS"wt", superfolder GFP | this study | https://benchling.com/s/seq-bMQCaaQ3rp9asp4FWGPm |
| pRBS"m1"-sfGFP | pACTet | RBS"m1", superfolder GFP | this study | https://benchling.com/s/seq-tUsA4giVksvanrJCPh0M |
| pRBS"m2"-sfGFP | pACTet | RBS"m2", superfolder GFP | this study | https://benchling.com/s/seq-PyKedE22cmYjZ1AbR5BN |
| pRBS"m3"-sfGFP | pACTet | RBS"m3", superfolder GFP | this study | https://benchling.com/s/seq-djklao9ARyOGmWDOucwH |
| pRBS"m4"-sfGFP | pACTet | RBS"m4", superfolder GFP | this study | https://benchling.com/s/seq-Z62DXUYRDRfFhiWt46dz |
